# Plant and prokaryotic TIR domains generate distinct cyclic ADPR NADase products

**DOI:** 10.1101/2022.09.19.508568

**Authors:** Adam M. Bayless, Sisi Chen, Sam C. Ogden, Xiaoyan Xu, John D. Sidda, Mohammad K. Manik, Sulin Li, Bostjan Kobe, Thomas Ve, Lijiang Song, Murray Grant, Li Wan, Marc T. Nishimura

## Abstract

Toll/interleukin-1 receptor (TIR) domain proteins function in cell death and immunity. In plants and bacteria, TIR domains are enzymes that produce isomers of cyclic ADPR (cADPR) as putative immune signaling molecules. The identity and functional conservation of cADPR isomer signals is unclear. A previous report found that a plant TIR could cross-activate the prokaryotic Thoeris TIR-immune system, suggesting the conservation of plant and prokaryotic TIR-immune signals. Here, we generate auto-active Thoeris TIRs and test the converse hypothesis: do prokaryotic Thoeris TIRs also cross-activate plant TIR-immunity? Using *in planta* and *in vitro* assays, we find that Thoeris and plant TIRs generate overlapping sets of cADPR isomers, and further clarify how plant and Thoeris TIRs activate the Thoeris system via producing 3’cADPR. This study demonstrates that the TIR-signaling requirements for plant and prokaryotic immune systems are distinct and that TIRs across kingdoms generate a diversity of small molecule products.

## Introduction

Globally, plant pathogens are estimated to diminish crop yields by over 15% each year and are a major threat to food security (Savary et al., 2012; Ficke et al., 2018). Understanding the mechanistic details of the plant immune system is a critical requirement for rationally engineering disease resistance. Unlike animals, plants do not possess adaptive immune systems and must encode expansive repertoires of cell-autonomous innate immune receptors to defend against pathogens. Toll/interleukin-1 receptor (TIR)-domains are encoded by plants, animals, and prokaryotes, and typically function in cell death and innate immune pathways (Nimma et al., 2017; Bayless and Nishimura, 2020; Essuman et al., 2022; Lapin et al., 2022). The TIR-domains of animal TLRs (Toll-like receptors) transduce immune signals via direct protein-protein interactions (Nimma et al., 2017). The discovery that the human TIR protein, SARM1, executes axonal degeneration via NAD^+^-hydrolase activity was pivotal to understanding TIR-immunity in plants and prokaryotes (Gerdts et al., 2015; Essuman et al., 2017; Essuman et al., 2022). Indeed, TIR-immune proteins of both plants and prokaryotes are now known to be enzymes which consume and/or modify nucleotides (including NAD^+^) or nucleic acids, and this enzymatic function is required for immune signaling across the tree of life (Horsefield et al., 2019; Wan et al., 2019; Essuman et al., 2022; Huang et al., 2022; Jia et al., 2022; Manik et al., 2022; Yu et al., 2022). The number and type of identified small molecules produced by enzymatic TIRs is expanding rapidly (Eastman et al., 2022a; Essuman et al., 2022; Lapin et al., 2022). Recently, it was reported that the TIR-immune signals of plants and of a prokaryotic anti-phage immune system, Thoeris, might be conserved (Ofir et al., 2021). The identity of the ThsB-TIR-produced Thoeris immune signal is unknown, and it is unclear if this prokaryotic immune signal might cross-activate plant TIR-immune pathways. Deciphering the identity and immune outputs of TIR-generated metabolites is key to understanding how to enhance plant TIR responses.

Plants initially sense potential pathogens via extracellular receptors that recognize conserved microbe-associated molecular patterns, or MAMPs (Jones et al., 2016). Upon binding MAMPs, these receptors trigger intracellular signaling cascades that activate a basal immune response known as PTI (pattern-triggered immunity) (Jones et al., 2016). However, adapted pathogens can disarm host PTI-responses via delivering virulence factors, or ‘effectors’, which manipulate host defense responses and/or physiology (Wang et al., 2022). Accordingly, plants have evolved a second layer of intracellular disease resistance proteins known as ‘R’ proteins, which recognize effectors or their activities and signal a rapid immune response termed ETI (effector-triggered immunity) (Jones et al., 2016). ETI often results in host cell death (the hypersensitive response), and plant ‘R-proteins’ generally contain N-terminal TIR or CC-domains (coiled coil) coupled to an NB-LRR (NLR) chassis (Jones et al., 2016). The C-terminal NLR domain confers effector recognition and provides an oligomerization chassis which promotes N-terminal TIR or CC-domain activation and immune signaling (Jones et al., 2016). TIR-NLR proteins are encoded by dicots, gymnosperms, and even single cellular algae (Shao et al., 2019; Bayless and Nishimura, 2020; Lapin et al., 2022). Curiously, monocots encode TIR-only (but not TNL) proteins which can cross-activate dicot TIR-immune pathways, although potential TIR-immunity within monocots remains less well characterized (Wan et al., 2019).

Like the immune TIRs of plants, the prokaryotic ThoerisB (ThsB) TIR generates immune signals using enzymatic activities (Ofir et al., 2021). However, Thoeris defense requires only a single downstream mediator/executioner to initiate cell death – the sirtuin2-type (SIR2) NADase, ThsA. Upon binding unknown ThsB-derived immune signals via a C-terminal SLOG-domain, ThsA causes host cell death by rapidly depleting cellular NAD^+^, thereby halting phage replication (Ofir et al., 2021). Apart from roles in immunity, certain microbial TIR-domain NADases have even been co-opted as virulence factors (Coronas-Serna et al., 2020; Eastman et al., 2022b). By contrast, plant TIR-immune signals must be relayed by several mediators: EDS1-family members (*Enhanced disease susceptibility 1*), and subsequently, the helper NLRs, NRG1 or ADR1 (N Requirement Gene 1, Activated Disease Resistance) (Bayless and Nishimura, 2020; Essuman et al., 2022). EDS1 is a lipase-like protein which forms exclusive heterodimers with *EDS1*-family members PAD4 (*Phytoalexin deficient 4*) or SAG101 (*Senescence-associated gene 101*), and these heterodimers were recently shown to bind the TIR-generated signals ADPr-ATP and pRib-AMP (Essuman et al., 2022; Huang et al., 2022; Jia et al., 2022; Lapin et al., 2022). Upon interacting with EDS1-heterodimers, the NRG1 or ADR1 helper NLRs oligomerize into Ca^2+^-permeable pores and transduce the initial TIR-immune signal into hypersensitive-cell death and/or transcriptional defense programs (El Kasmi, 2021; Jacob et al., 2021; Kim et al., 2022; Tian and Li, 2022).

Certain mechanistic features of enzymatic TIRs are conserved even across very distant phyla (Essuman et al., 2018; Wan et al., 2019; Essuman et al., 2022). For instance, all examined prokaryotic and eukaryotic TIRs require a conserved glutamate (E) residue for catalysis (Essuman et al., 2017; Horsefield et al., 2019; Ofir et al., 2021; Essuman et al., 2022; Manik et al., 2022). Additionally, enzymatic TIRs contain a flexible loop, termed the BB-loop, which lays over the catalytic pocket and has been proposed to regulate substrate access (Ma et al., 2020; Martin et al., 2020; Manik et al., 2022; Shi et al., 2022). Enzymatic TIR-domains also require self-association via TIR-TIR interfaces to engage in catalysis (Horsefield et al., 2019; Wan et al., 2019; Manik et al., 2022; Shi et al., 2022). Cryo-EM studies have provided key insights into how oligomerization and self-association promote the activation of plant TIR-NLRs, as well as CC-NLRs (Wang et al., 2019a; Wang et al., 2019b; Ma et al., 2020; Martin et al., 2020). For example, the structure of the pentameric ZAR1 ‘resistosome’ reveals that pathogen effectors induce the assembly of CC-NLRs into ring-shaped Ca^2+^-ion channels (Wang et al., 2019a; Bi et al., 2021). Similarly, the structures of the activated TIR-NLRs, RPP1 (*Recognition of Peronospera parasitica 1*) and Roq1 (*Recognition of XopQ1*), indicate that effector activities also induce TIR-NLRs to assemble into tetramers, thereby engaging their enzymatic cores (Ma et al., 2020; Martin et al., 2020). The crystal structure of the Thoeris ThsB TIR-protein reveals a core-TIR-domain followed by a small C-terminal β-sheet domain; the structure also suggests that ThsB could form dimers (Ka et al., 2020). Bacteriophage triggers the TIR-NADase activity of ThsB, although the mechanistic details of activation and potential ThsB-ThsB oligomerization requirements are unknown (Ka et al., 2020; Ofir et al., 2021). Plants also encode TIR-domains which lack NLR architecture, including TIR-NB, TIR-only proteins, and XTNX proteins (Nandety et al., 2013; Nishimura et al., 2017; Johanndrees et al., 2021). A recent study found that a plant TIR-only protein, RBA1, can self-associate and perform enzymatic functions as linear filaments on nucleic acids (Yu et al., 2022). The atypical XTNX TIR-architecture may activate cell death independently of the TIR-pathway mediator, *EDS1* (Johanndrees et al., 2021).

As noted, plant and prokaryotic immune TIRs, as well as human SARM1, require enzymatic activity for their functions in immunity and axonal cell death, respectively. SARM1 NADase activity produces ADPR and canonical cyclic ADPR (cADPR), which are both secondary messengers that can trigger intracellular Ca^2+^-signaling cascades within animal cells (Fliegert et al., 2020; Eastman et al., 2022a; Li et al., 2022). Intriguingly, the TIR-domains of plants and prokaryotes produce non-canonical cADPR-isomers from NAD^+^, such as 3’cADPR or 2’cADPR (Essuman et al., 2018; Wan et al., 2019; Essuman et al., 2022; Manik et al., 2022; Weagley et al., 2022). 3’cADPR is the NADase product of the HopAM1 TIR-protein - a virulence factor encoded by the plant pathogen *Pseudomonas syringae DC3000* (Eastman et al., 2022b). Initially, 2’cADPR was known as ‘variant cADPR’ (v-cADPR) given its unknown structure but identical mass to canonical cADPR (MW: 541 Da) (Essuman et al., 2018; Wan et al., 2019). While v-cADPR (2’cADPR) is a biomarker of plant TIR-pathway activation by pathogens, a later study reported that v-cADPR-generation by a prokaryotic TIR, AbTir (from human pathogen *Acinetobacter baumanii*), was insufficient to stimulate *EDS1-*mediated plant hypersensitive (HR) cell death (Duxbury et al., 2020). Potential conservation among plant and prokaryotic TIR-immune signals remains unclear (Ofir et al., 2021; Leavitt et al., 2022). Here, we examined whether the NADase-derived signals from the Thoeris immune TIR, ThsB, are functionally compatible with plant TIR-pathways.

## Results

### Prokaryotic TIR NADases cause cytotoxicity independent of the plant TIR-signal mediator, EDS1

Plant TIR-enzymatic activities are required to signal *EDS1-*mediated hypersensitive cell death (HR) (Horsefield et al., 2019; Wan et al., 2019). As plant BdTIR cross-activated ThsA, the mediator of Thoeris immunity (Ofir et al., 2021), we tested if the ThoerisB (ThsB) immune TIR (of *B. cereus* MSX D-12) could generate signals that cross-activated *EDS1 in planta* (see schematic diagram in Fig. 1*A*). Accordingly, we synthesized and expressed codon-optimized ThsB in the plant *Nicotiana benthamiana* (*Nb*) and monitored *EDS1-*mediated HR (hypersensitive cell death) (Fig. 1*B*, *C*). We expressed plant BdTIR as a positive control for HR, and the human TIR NADase, SARM1, as an *EDS1-*independent cell death inducer (via NAD^+^-depletion) (Wan et al., 2019). Additionally, we expressed the core TIR-domain of AbTir (core-TIR denoted as ‘AbTIR’), a non-immune TIR from the pathogenic bacterium *Acinetobacter baumannii*, which generates 2’cADPR (Duxbury et al., 2020; Manik et al., 2022). BdTIR expression triggered HR in wild-type (WT) *Nb*, but not in plants lacking *EDS1* (*Nb eds1*^−/−^), while SARM1-TIR elicited cell death in both (Fig. 1*B*, *C*). Similar to SARM1-TIR, AbTIR triggered chlorotic cell death in both WT and *eds1^−/−^* plants, while ThsB caused no apparent phenotype (Fig. 1*B*, *C*). We also assayed another v-cADPR-producing TIR-domain from the Archaea *Methanobrevibacter olleyae* (TcpO-TIR), which likewise caused mild chlorosis independent of *EDS1* (Fig. S1) (Essuman et al., 2018; Wan et al., 2019). Chlorotic cell death by AbTIR or TcpO-TIR required the conserved TIR-domain catalytic glutamate (E), indicating that while these TIRs are enzymatically active, neither produces *EDS1*-activating signals (Fig. 1*B*, *C*). We also observed that elevating BdTIR expression (via adding Omega translational enhancers (Gallie, 2002)) could similarly cause a slow chlorotic cell death independent of *EDS1* (Fig. S1). AbTIR, TcpO-TIR, and ThsB were phenotyped without epitope tags, while N-HA tags were used to confirm expression via immunoblot (Fig. 1*C*, *D*). Structural models indicating the conserved TIR-domain structure and catalytic glutamate (E) of AbTir and TcpO are provided in Figure S1*A* and *B*. Like plant TNL immune receptors, ThsB NADase activities are triggered by pathogens (*i.e*., bacteriophages) and the lack of an apparent phenotype suggested that ThsB might be inactive in the absence of this trigger (Ka et al., 2020; Ofir et al., 2021).

**Figure 1.**
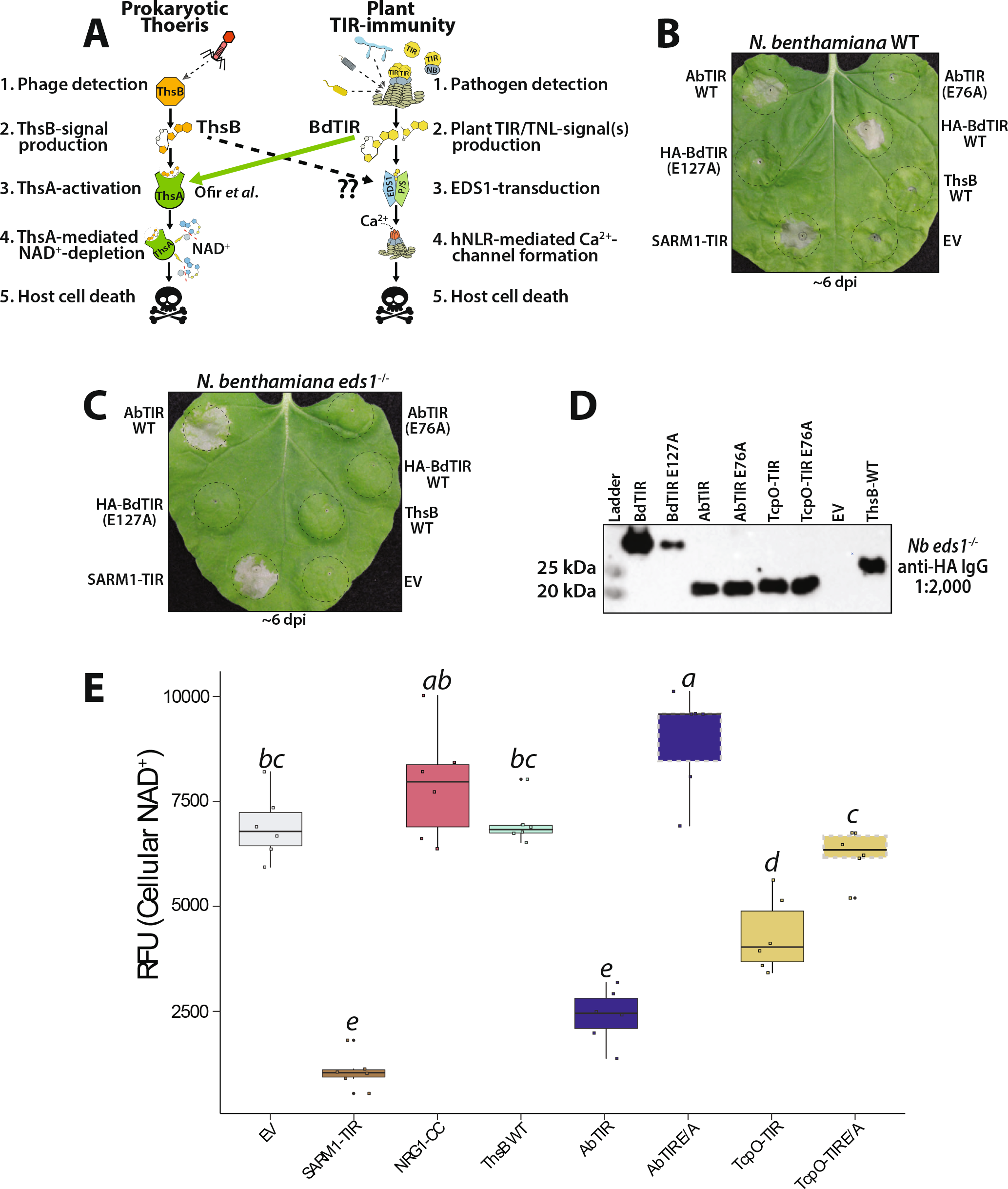
Prokaryotic TIRs can trigger in planta cytotoxicity independently of EDS1. (*A*) Schematic diagram of prokaryotic Thoeris immunity and plant TIR-immune signaling. In both immune systems, TIRs sense pathogens and signal immune outputs via NADase activities. BdTIR stimulates Thoeris-outputs; does ThsB stimulate *EDS1*-outputs? (*B*, *C*) *Nb* WT or *eds1*^−/−^ leaves shown approximately 6 days post *Agro*-infiltration with constructs delivering AbTIR, HA-BdTIR, ThsB, or EV (empty vector control; 35s-GFP). E/A refers to TIRs containing alanine substitutions at the conserved catalytic glutamate (E) residue. (*D*) Anti-HA immunoblot of N-HA tagged BdTIR, AbTIR, ThsB or TcpO-TIR proteins harvested from *Nb eds1*^−/−^ leaves at ~40 hours post-infiltration (hpi). (*E*) Fluorescent NAD^+^-detection assay in *Nb eds1*^−/−^ leaves expressing different TIR constructs. NAD^+^-assays were performed at 40 hpi. NRG1-CC is the coiled-coil domain of the CNL protein, N Requirement Gene 1; SARM1: Sterile alpha and TIR-motif containing 1. Catalytic glutamate (E residue) mutants of TIRs outlined in dotied grey boxes. Statistical analyses: One-way ANOVA and Turkey HSD with CLD (compact letier display) of significance classes. Over-lapping letiers are ns (non-significant) difference (*p*>.05) while separate letier class indicates *p*<.05 or better.

SARM1-mediated cell death in neurons and plants correlates with a strong depletion of NAD^+^ (Essuman et al., 2017; Horsefield et al., 2019; Wan et al., 2019). Thus, we used a fluorescence-based NAD^+^-assay to determine if AbTIR, TcpO-TIR or ThsB depleted NAD^+^ *in planta* (Fig. 1*E*). SARM1-TIR was included as a positive control for NAD^+^-depletion (Fig. 1*E*). AbTIR and TcpO-TIR both reduced cellular NAD^+^, suggesting that the *EDS1*-independent toxicity in Fig. 1*B* may be explained by perturbation of NAD^+^-homeostasis, similar to SARM1-TIR. NRG1-CC causes rapid cell death via Ca^2+^-channel formation and serves as a control to indicate that plant cell death *per se* does not drive NAD^+^-depletion (Fig. 1*E*). As above, ThsB elicited no apparent NAD^+^ depletion, in spite of abundant protein accumulation (Fig. 1*D*). To try to find an active ThsB allele, we assayed 11 additional ThsB orthologs encoded by other bacterial species; none caused apparent phenotypes (Fig. S2). Altogether, these results suggest that while v-cADPR-producing AbTIR and TcpO-TIR do not trigger *EDS1*-dependent immunity, highly active TIR NADases can be cytotoxic via cellular NAD^+^-depletion. Neither TcpO or AbTir have known roles in host immune signaling. Thus, we refocused on generating ThsB enzymatic activity *in planta*, to assess the conservation of plant and prokaryotic TIR-immune signals.

### Replacement of a ThsB loop region promotes auto-activity and stimulation of the Thoeris-mediator, ThsA

Thoeris immunity (*B. cereus* MSX D12) is initiated by phage detection, indicating that ThsB signaling is inactive prior to infection (Ofir et al., 2021). Hence, we hypothesized that an auto-active ThsB NADase might be created by removing or modifying negative regulatory regions within ThsB. An auto-active ThsB should hydrolyze NAD^+^ and generate cADPR-isomers which stimulate the Thoeris partner, ThsA. Also, if the TIR-immune signals of plants and the Thoeris system are conserved, an auto-active ThsB might stimulate *EDS1*-mediated HR (see Fig. 1*A* model).

A previous study by Ka et al. determined the crystal structures of ThsB, and the mediator, ThsA (Ka et al., 2020). To gain insights into potential ThsB regulatory regions, we docked ThsB onto the structure of an activated-state plant TIR-NLR, RPP1 (*Recognition of Peronospora parasitica 1*) (Ka et al., 2020; Ma et al., 2020) (Fig. 2*A*). The activated structures of RPP1 and ROQ1 TIR-domains indicate that movement of a loop (the “BB-loop”) near the catalytic glutamate may be important for allowing substrate access and/or catalysis (Ma et al., 2020) (Fig. 2*A*). We noted that ThsB also has a putative BB-loop in this position, although it is not resolved in the ThsB crystal structure. Unlike plant TIR-domains, ThsB contains a large C-terminal β-sheet following the TIR-domain (Fig. 2*A*). Thus, we hypothesized that this loop near the catalytic pocket, and/or the C-terminal β-sheet might affect ThsB NADase activation. Accordingly, we generated and examined five different ThsB variants for auto-activity: a modified ‘loop’-region, and four progressive deletions of the ‘β-sheet’-domain (Fig. 2, S3). Briefly, the ‘loop’ variant substituted loop residues with glycines, while each C-terminal variant removed the denoted residues and added a short glycine-linker. These five ThsB variants were named: ‘Auto’ (loop region), ‘Core-TIR’, ‘120-165’, ‘145-163’, or ‘152-163’. Fig. S3 maps each of these modifications onto the ThsB crystal structure.

**Figure 2.**
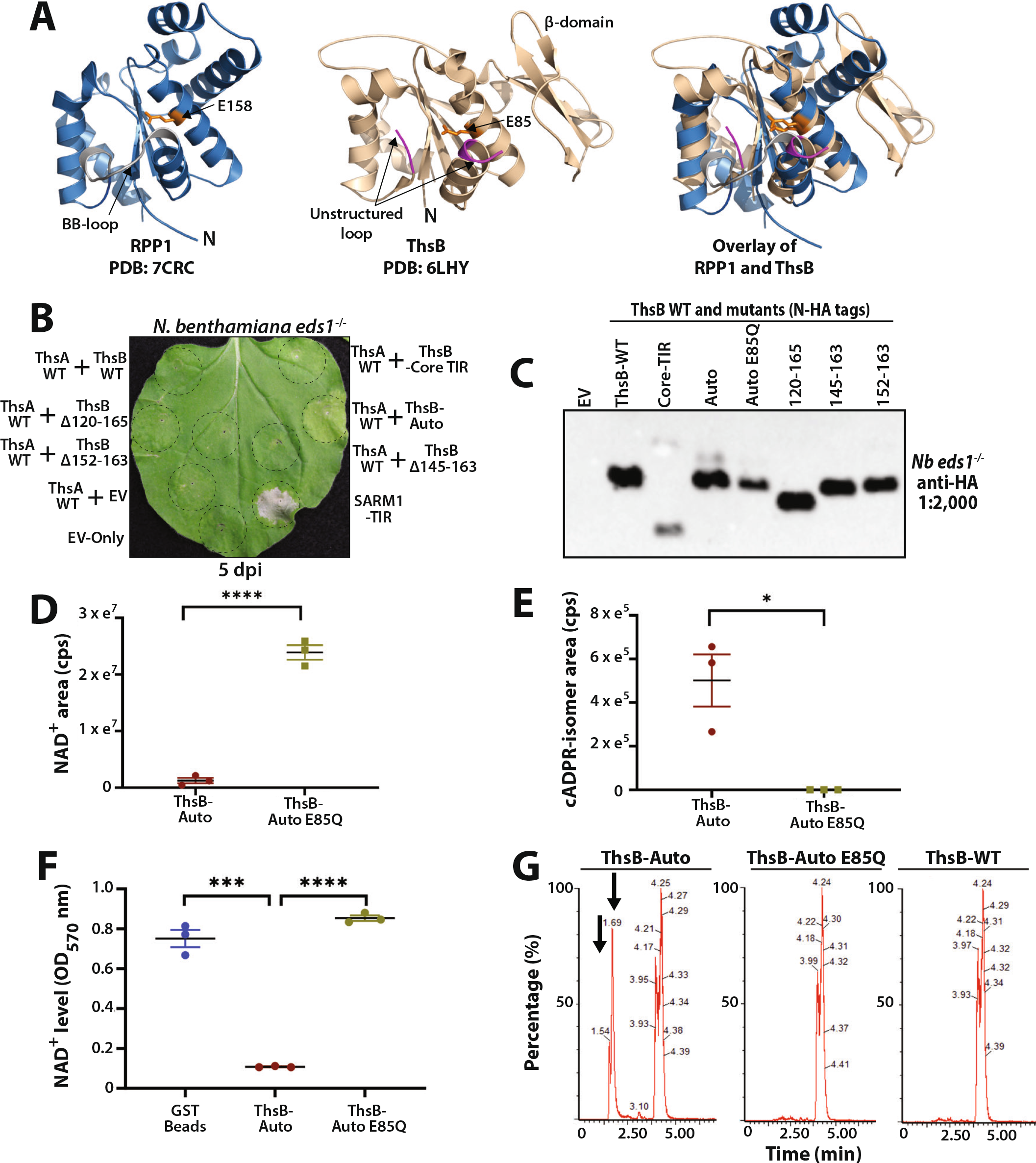
Mutagenesis of a ThsB loop region promotes NADase auto-activity, cADPR-isomer production, and stimulation of ThsA-mediated cytotoxicity. (*A*) (Leti) Cryo-EM structure of activated state TIR-domain from plant TNL, RPP1 (PDB: 7CRC). (Center) Crystal structure ThsB (PDB: 6LHY). (Right) Overlay of activated RPP1-TIR with ThsB. The RPP1 BB-loop is colored grey, ThsB loop region shown in purple, and catalytic glutamates shown in orange. N: N-terminus. (*B*) *Nb eds1*^−/−^ leaves ~5 days post *Agro*-infiltration (dpi) with constructs co-expressing ThsA with either WT ThsB, ThsB-mutants, or EV (empty vector control; 35S-GFP). ThsA and ThsB were co-expressed at an individual OD of 0.40. Positive and negative control SARM1 or EV, respectively, expressed individually at OD 0.80. (*C*) Anti-HA immunoblot of N-HA tagged ThsB variants transiently expressed in *Nb eds1*^−/−^ leaves and harvested ~40 hpi. (*D*, *E*) *In vitro* NAD^+^-consumption by recombinant ThsB-Auto or ThsB-Auto E85Q (catalytic mutant), and detection of cADPR-isomer via LC-MS. LC-MS traces for cADPR-isomers (MW 542) in *Nb eds1*^−/−^ leaves transiently expressing ThsB-Auto or ThsB-Auto E85Q; leaves sampled ~40 hpi. (*G*) LC-MS traces for cADPR-isomers (MW 542) in *Nb eds1*^−/−^ leaves transiently expressing ThsB-Auto, ThsB-Auto E85Q, or WT ThsB; leaves sampled ~40 hpi. Arrows denote ThsB-Auto produced cADPR-isomers. Statistical analyses: One-way ANOVA and Turkey HSD. ns: not significant, \**p*<0.05, \*\**p*<0.01, \*\*\**p*<0.005, \*\*\*\**p*<0.0005.

Because the TIR-domains of AbTir and SARM1 deplete cellular NAD^+^ and cause chlorotic cell death independent of *EDS1* (Fig. 1*B*), we reasoned that ThsB-activation of ThsA, a SIR2-type NADase, might cause similar chlorosis and cytotoxicity. Thus, we screened each ThsB variant for auto-activity by co-expressing it with ThsA and monitoring for chlorosis (Fig. 2*B*). ThsA co-expression with WT ThsB, or any C-terminal deletion, did not trigger chlorosis (Fig. 2*B*). An anti-HA immunoblot confirmed accumulation for all ThsB variants (Fig. 2*C*). When ThsA was co-expressed with the ThsB-loop deletion, termed ‘Auto’, a mild chlorotic cell death appeared ~4-5 dpi (Fig. 2*B*). To attempt to increase the activity of ThsB-Auto, we generated additional mutants in the BB-loop (Fig. S3*C*, *D*). While the new BB loop mutants were also auto-active, neither had appreciably stronger phenotypes than the original ThsB-Auto mutant. We also examined the effects of similarly altering the BB-loop of plant TIR proteins, however, we found that this caused a loss of *EDS1-*signaling function (Fig. S3*E*, *F*, *G*).

We next determined if ThsB-Auto produced cADPR-isomers from NAD^+^ (Fig. 2*D*-*G*). Using recombinant ThsB-Auto protein, we performed *in vitro* NADase assays and LC-MS, and detected the consumption of NAD^+^ and cADPR-isomer generation (Fig. 2*D*, *E*). Furthermore, the NADase products of ThsB-Auto could stimulate NAD^+^-consumption by ThsA *in vitro* (Fig. 2*F*). We also used LC-MS and detected cADPR-isomer production by ThsB-Auto *in planta* (Fig. 2*G*). cADPR-isomer generation by ThsB required the catalytic glutamate (E85), as did *in vitro* stimulation of ThsA (Fig. 2*D*-*G*). Together, these findings indicate that the TIR NADase functions of ThsB can be activated by modifying a portion of its BB-loop and suggest that this loop may play a part in regulating Thoeris-signaling. Importantly, the NAD^+^-breakdown products from ThsB-Auto stimulated ThsA activity *in vitro*, in addition to stimulating *in planta* cytotoxicity (Fig. 2*B*).

### ThsB-Auto requires the TIR-domain catalytic glutamate to stimulate ThsA-mediated NAD^+^-depletion and cytotoxicity

During Thoeris immunity, stimulated ThsA drives host cell death via NAD^+^-depletion and restricts phage replication (Ofir et al., 2021). To validate ThsA-stimulation by ThsB-Auto *in planta*, we expressed ThsA and ThsB alone, or together, and examined cellular NAD^+^ levels and cytotoxicity in *Nb eds1*^−/−^ plants (Fig. 3*A*-*E*). As controls, we included previously described ThsA N112A, which lacks SIR2-NADase activity, and ThsA R371A, which is reportedly impaired in binding activating cADPR-isomer (Ka et al., 2020; Ofir et al., 2021; Manik et al., 2022). Figure S4 maps N112 and R371 onto the ThsA crystal structure (Ka et al., 2020). When expressed alone, neither WT ThsB, ThsB-Auto, or ThsB-Auto E85Q, diminished NAD^+^ or caused cytotoxicity similar to positive control SARM1-TIR (Fig. 3*A*, *B*). ThsA alone has been reported to have some constitutive, but weak background NADase activity *in vitro* (Ka et al., 2020; Ofir et al., 2021; Manik et al., 2022). Consistent with this, ThsA mildly reduced NAD^+^ relative to empty vector or NADase-null ThsA N112A controls *in planta* (Fig. 3*B*). Importantly, NAD^+^-consumption by unstimulated ThsA was not enough to trigger qualitative or quantitative cytotoxicity (Fig. 3*A*, *C*, *E*). Surprisingly, ThsA R371A displayed enhanced NAD^+^-depletion compared to WT ThsA, and accordingly, caused macroscopic cell death like SARM1-TIR (Fig. 3*A*). This was unexpected, and so we further examined whether disruption of the ThsA SLOG-motif enhanced background NADase activities (Fig. S4). We generated three additional SLOG mutations (E403A, K388A, K388E), and all had enhanced NADase activity relative to WT ThsA and did not require stimulation by ThsB-Auto (Fig. S4*B*, *C*). Regardless, these SLOG-mutants illustrate that enhancing ThsA NADase activities can sharply deplete cellular NAD^+^ and trigger macroscopic cell death. Further, while the SLOG-motif facilitates cADPR-isomer binding and NADase activation, it also apparently regulates ThsA activation in the absence of signal. Collectively, these findings demonstrate that neither ThsB-Auto, nor unstimulated WT ThsA deplete cellular NAD^+^ to an extent that is cytotoxic.

**Figure 3.**
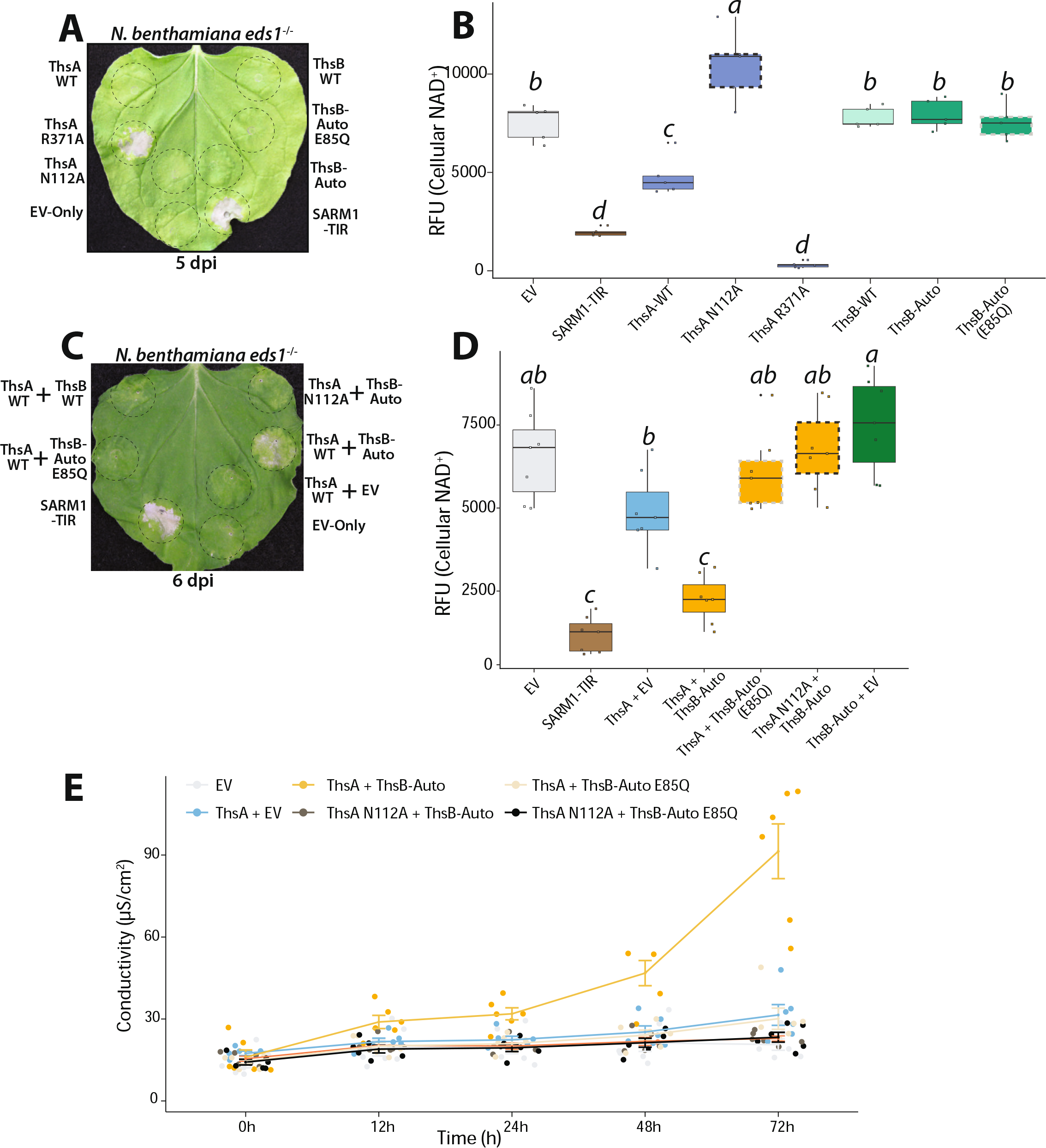
ThsB-Auto requires TIR-NADase functions to stimulate ThsA-mediated NAD^+^-depletion and cytotoxicity. (*A*) *Nb eds1*^−/−^ leaves ~5 dpi with constructs expressing WT ThsA or ThsB, ThsA and ThsB variants, or SARM1 or EV (35S-GFP) controls. ThsA N112A lacks SIR2-type NADase activity; ThsA R371A has an altered SLOG-motif, and ThsB E85Q lacks TIR-domain catalytic activity. Constructs expressed at OD 0.80. (*B*) Fluorescent NAD^+^-detection assay in *Nb eds1*^−/−^ leaves expressing different ThsA or ThsB constructs. NADase-assays performed 40 hpi. (*C*) *Nb eds1*^−/−^ leaves co-expressing ThsA or ThsA N112A with ThsB-Auto, ThsB-Auto E/Q, or EV, shown ~6 dpi. ThsA and ThsB co-expressed at OD 0.40; positive and negative control SARM1 and EV expressed at OD 0.80. Catalytic TIR (E/A) mutants outlined in dotied grey; ThsA N112A outlined in dotied black. (*D*) Fluorescent NAD^+^-detection assay in *Nb eds1*^−/−^ leaves co-expressing ThsA and ThsB combinations. NAD^+^-assays performed 40 hpi. Catalytic glutamate mutants are outlined in dotied grey; inactive ThsA N112A are outlined in dotied black. (*E)* Ion leakage assay in *Nb eds1*^−/−^ leaves co-expressing ThsA and ThsB combinations. Leaf discs were collected ~72 hpi. Statistical analyses: One-way ANOVA and Turkey HSD with compact letier display of significance classes. Over-lapping letiers are ns (non-significant) difference (*p*>.05) while separate letier class indicates *p*<.05 or better.

We then validated that ThsA-stimulation by ThsB-Auto required the TIR-domain likely catalytic glutamate (E) residue (Fig. 3*C*, *D*, *E*). As in Figure 2, we co-expressed ThsA WT with ThsB WT, ThsB-Auto, or ThsB-Auto E85Q, and monitored for chlorotic cell death in *Nb eds1*^−/−^ plants (Fig. 3*C*). ThsA co-expression with ThsB-Auto elicited chlorosis, while co-expression with either ThsB WT or ThsB-Auto E85Q had no effect. Similarly, ThsB-Auto co-expression with NADase null ThsA N112A did not trigger cell death (Fig. 3*C*). ThsA and ThsB-Auto co-expression also sharply reduced cellular NAD^+^, as compared to control pairings of ThsB-Auto E85Q or ThsA N112A (Fig. 3*D*). To quantitatively assess cytotoxicity from ThsA-stimulation by ThsB-Auto, we performed ion leakage assays on leaf discs (Fig. 3*E*). Briefly, during the onset of plant cell death, cellular ions leak into solution due to loss of membrane integrity. Consistent with the macroscopic cell death and NAD^+^-depletion observed in Fig. 3*C* and *D*, we recorded significant increases in ion conductivity when ThsA was co-expressed with ThsB-Auto, but not with ThsB-Auto E85Q or ThsA N112A loss-of-function controls (Fig. 3*E*). Altogether, these findings indicate that the catalytic activities of ThsB-Auto are required to stimulate ThsA-mediated NAD^+^-depletion and cell death. Furthermore, the functional inputs and outputs of the prokaryotic Thoeris system can be reconstructed *in planta*.

### A plant TIR, BdTIR, cross-activates ThsA, but ThsB-Auto does not cross-activate EDS1-mediated cell death

The plant TIR BdTIR has been reported to activate ThsA, hinting that TIR-signals from plant and bacterial immune systems might be conserved (Ofir et al., 2021). After validating ThsB-Auto stimulation of ThsA *in planta*, we tested the reverse hypothesis: does ThsB-Auto stimulate HR-outputs by the plant TIR-mediator, *EDS1*? Accordingly, we expressed BdTIR and ThsB-Auto in WT *Nb* and monitored HR-cell death (Fig. 4*A*). As in Figure 1*C*, BdTIR signaled *EDS1-*mediated cell death, however, no cell death phenotypes were observed from ThsB-Auto, ThsB WT, or any ThsB variant (Fig. 4*A*). The finding that ThsB-Auto stimulates ThsA but is insufficient to activate *EDS1* indicates that the signaling requirements for plant and Thoeris pathways are different. It further suggests that the enzymatic products of ThsB-Auto and plant TIRs are likely distinct.

**Figure 4.**
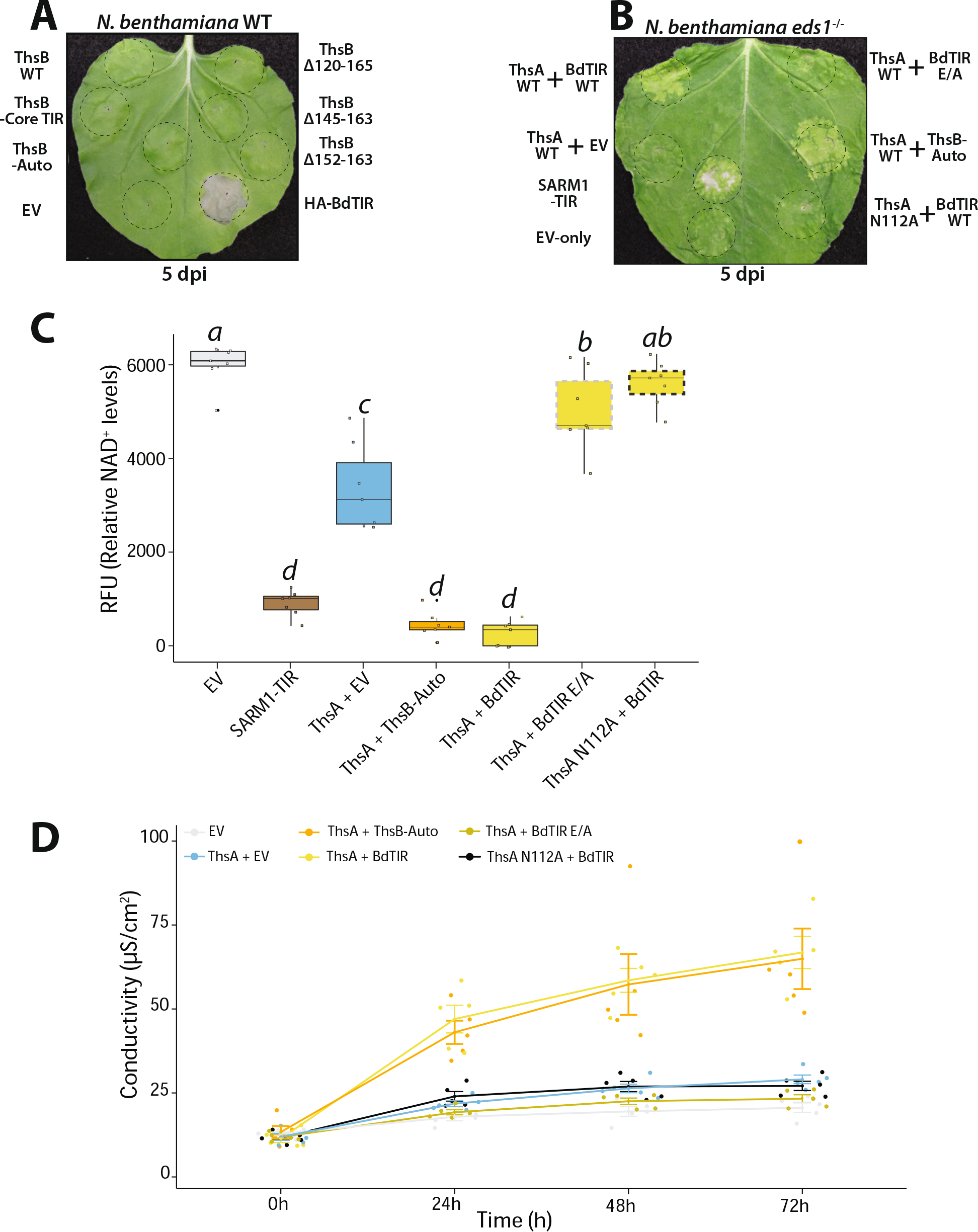
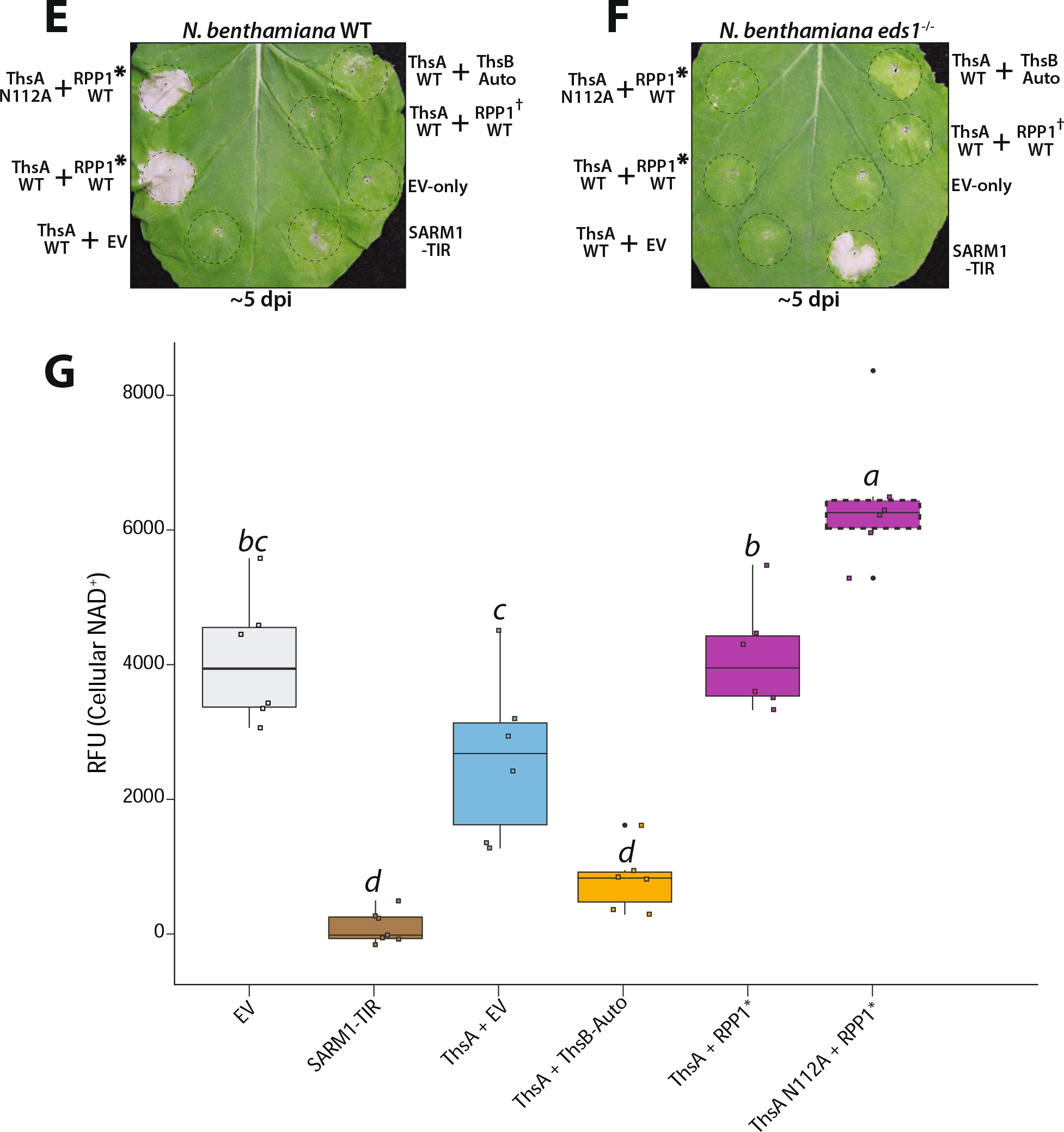
ThsB-Auto does not stimulate EDS1-mediated HR. BdTIR does stimulate ThsA, but the activated TNL, RPP1, does not. (*A*) *Nb* WT leaves ~5 days post *Agro*-infiltration with constructs expressing HA-BdTIR, ThsB WT, ThsB-Auto, or the previously noted ThsB C-terminal mutations. All constructs, including negative control EV, were expressed at OD 0.80. (*B*) *Nb eds1*^−/−^ leaves ~6 days post *Agro*-infiltration with constructs co-expressing ThsA or ThsA N112A with HA-BdTIR or BdTIR E/A. ThsA with ThsB-Auto and EV controls were also included. All co-expressions contained OD 0.40 of each construct, while SARM1 and EV positive and negative controls were at OD 0.80. BdTIR E/A lacks TIR-domain catalytic activity. (*C*) Fluorescent NAD^+^-detection assay in *Nb eds1*^−/−^ leaves co-expressing ThsA, HA-BdTIR or ThsB-Auto combinations, or positive and negative control SARM1 and EV. NAD^+^-assays performed 40 hpi. (*D*) Ion leakage assay in *Nb eds1*^−/−^ leaves co-expressing different ThsA and ThsB combinations. Leaf discs were collected ~72 hpi, and measurements recorded every 24 h for 3 days. (*E, F*) *Nb* WT or *eds1*^−/−^ leaves ~5 days post *Agro*-infiltration with constructs co-expressing ThsA, ThsB-Auto, or RPP1-TNL with cognate ATR1 effector. *denotes RPP1_WsB with activating ATR1-Emoy effector, while † has non-activating ATR1-Emwa. ThsA co-expressed at OD 0.40, RPP1/ATR1 expressed at OD 0.20 each. (*G*) Fluorescent NAD^+^-detection assay in *Nb eds1*^−/−^ leaves co-expressing noted ThsA, RPP1/ATR1, or ThsB-Auto combinations, as well as positive and negative control SARM1 and EV. NAD^+^-assays performed 40 hpi. Statistical analyses: One-way ANOVA and Turkey HSD. Over-lapping letiers are ns (non-significant) difference (*p*>.05) while separate letier class indicates *p*<.05 or better.

Given the above lack of *EDS1*-activation by ThsB-Auto, we verified that BdTIR cross-activates ThsA (Ofir et al., 2021) (Fig. 4*B*, *C*, *D*). As in Figure 3, we co-expressed ThsA with BdTIR or ThsB-Auto and examined NAD^+^-depletion and cell death (Fig. 4*B*, *C*, *D*). Like ThsB-Auto, BdTIR stimulated ThsA-mediated cytotoxicity and NAD^+^-consumption (Fig. 4*B*, *C*, *D*), confirming that signals from a plant TIR can also cross-active the Thoeris system *in planta* (Ofir et al., 2021).

### The dicot TIR resistance protein, RPP1, signals EDS1-HR but does not stimulate ThsA

Although BdTIR activates *EDS1-*mediated HR in the dicot plant, *Nicotiana benthamiana*, *BdTIR* has no demonstrated immune functions within the monocot, *Brachypodium distachyon*. Therefore, we tested if the TIR-NLR, RPP1 (<u>R</u>ecognition of *<u>P</u>eronospora <u>p</u>arasitica* <u>1</u>), from the dicot *Arabidopsis thaliana*, also stimulated ThsA (Fig. 4*E*, *F*). Accordingly, we expressed ThsA with ThsB-Auto, or RPP1 and cognate ATR1 effector (*<u>A. t</u>haliana* <u>R</u>ecognized <u>1</u>). While ATR1 activated RPP1 to signal HR, ATR1-activation of RPP1 was unsuccessful at stimulating ThsA-mediated chlorosis or NAD^+^-depletion (Fig. 4*E*, *F*). The incompatibility of ThsB-Auto to initiate *EDS1-*dependent cell death, along with the finding that activated RPP1 does activate *EDS1* but not ThsA, further suggests that the input signals for plant and prokaryotic TIR-systems are not conserved.

### ThsB-Auto and BdTIR produce different primary isomers of cADPR; ThsA is efficiently stimulated by 3’cADPR producing TIRs

Recently, Eastman et al. reported that the TIR-effector protein, HopAM1, produced a cADPR-variant unlike that made by AbTIR; these two variant cADPRs have been identified as 2’cADPR (AbTIR product) and 3’cADPR (HopAM1 product) (Eastman et al., 2022b; Manik et al., 2022). To determine which cADPR-isomer(s) ThsB-Auto produced, we compared the *in vitro* NADase products of ThsB-Auto, BdTIR, AbTIR, and HopAM1 using LC-MS (Fig. 5*A*). The major cADPR-isomer produced by ThsB-Auto had a unique retention time, unlike canonical cADPR-standard or any products from BdTIR, HopAM1 or AbTIR (Fig. 5*A*). ThsB-Auto also produced a second peak, aligning with 3’cADPR made by HopAM1 (Fig. 5*A*). Unexpectedly, BdTIR generated both 2’cADPR (major product) and 3’cADPR (minor product) while AbTIR only produced 2’cADPR (Fig. 5*A*). Quantification of 3’cADPR peaks is shown in Figure 5*B*, *C*. Importantly, these findings provide a testable prediction: ThsA is stimulated by TIRs that produce 3’cADPR, but not 2’cADPR.

**Figure 5.**
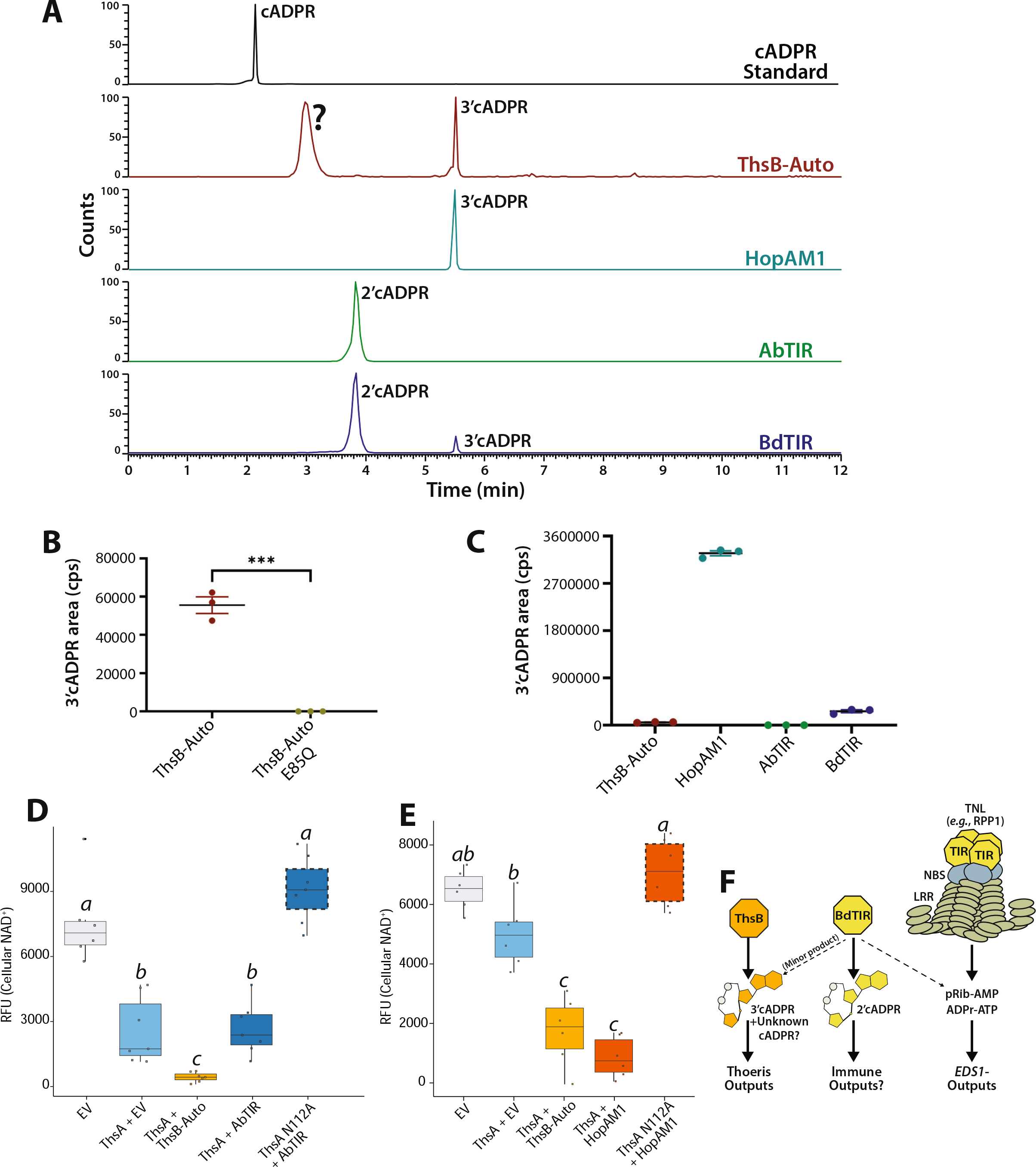
ThsB-Auto and BdTIR produce different cADPR-isomers as major products; ThsA is highly stimulated by 3’cADPR. (*A, B*) *In vitro* production of 3’cADPR by ThsB-Auto relative to E85Q, or HopAM1, AbTIR or BdTIR. (*C*) LC-MS traces of cADPR-isomers from *in vitro* NADase reactions of ThsB-Auto, HopAM1, AbTIR or BdTIR, relative to cADPR standard. (*D, E*) Fluorescent NAD^+^-detection assay in *Nb eds1*^−/−^ as previously described, except with AbTIR or HopAM1 co-expressed to stimulate ThsA. NAD^+^-assays performed at ~40 hpi. (*F*) Model of Thoeris and *EDS1*-pathway stimulation by signals from plant and prokaryotic TIRs. Statistical analyses: One-way ANOVA and Turkey HSD. Over-lapping letiers are ns (non-significant) difference (*p*>.05) while separate letier class indicates *p*<.05 or better.

Ofir et al. showed that ThsA was stimulated by cADPR-isomers from ThsB and BdTIR, but not canonical cADPR, while Manik et al. found that ThsA orthologs from *Enterococcus faecium* and *Streptococcus equi* were highly stimulated by 3’cADPR but not 2’cADPR (Ofir et al., 2021; Manik et al., 2022). Given that both BdTIR and ThsB-Auto make 3’cADPR and stimulate ThsA, we predicted that HopAM1 (3’cADPR producer) would stimulate ThsA while AbTIR (2’cADPR) would not. We therefore co-expressed ThsA with AbTIR, or HopAM1, and assayed cell death and NAD^+^-depletion as before in *Nb eds1^−/−^* plants (Fig. 5*D*, *E*, S5, S6). AbTIR co-expression did not enhance ThsA-mediated cytotoxicity or NAD^+^-depletion relative to ThsB-Auto (Fig. 5*D*, S5). Because HopAM1 is cytotoxic to *Nb* when highly expressed, we first titrated the HopAM1 delivery and found that OD_600_ < 0.010 did not deplete NAD^+^ or trigger cell death (SI 5). When co-expressed at OD 0.005, HopAM1 stimulated NAD^+^-depletion by ThsA similar to that promoted by ThsB-Auto (Fig. 5*E*).

ThsB-Auto stimulates ThsA and makes 3’cADPR, as well as an undefined cADPR isomer (Fig. 5*A*). Whether other ThsB alleles might also produce this uncharacterized isomer was unclear, however. Therefore, we examined if any of the previously screened ThsB-orthologs (see Fig. S2) could stimulate ThsA, and if they also produced this atypical isomer (Fig. S7). The TIR-domain of a second ThsB (ThsB2-TIR) encoded by *B. cereus* (MSX D-12) signaled ThsA-mediated cell death in *N. benthamiana*, while no other tested ortholog did (Fig. S7) (Doron et al., 2018). Similarly, ThsB2-TIR promoted ThsA-mediated NAD^+^-depletion, while LC-MS analysis revealed that ThsB2 produced 3’cADPR, but not the undefined isomer made by ThsB-Auto (Fig. S7). ThsB2-TIR more effectively stimulated ThsA-mediated cell death than the full-length ThsB2 protein, indicating that the N-terminus may impact NADase activities (Fig. S7). ThsB (192 aa) and ThsB2 (193 aa) differ in length by just 1 residue, however, their overall amino acid identity is low (<20%). AlphaFold predictions suggest that ThsB and ThsB2 do not share structural homologies outside the core-TIR domain (Fig. S7). Notably, ThsB2-TIR did not require BB-loop modifications to stimulate ThsA, suggesting that the NADase activities of ThsB and ThsB2 could be regulated differently.

These findings identify 3’cADPR as an efficient trigger for ThsA and suggest that ThsB-Auto produced 3’cADPR is the activating signal for the Thoeris system. We further explain the cross-stimulation of the Thoeris system by BdTIR: BdTIR generates multiple isomers of cADPR, including the 3’cADPR made by ThsB-Auto. The model in Figure 5*F* summarizes these results, while Fig. S8 lists the known enzymatic products of each TIR and their *EDS1*/ThsA-stimulation phenotypes.

## Discussion

Plant and prokaryotic TIRs use enzymatic activities to produce immune signals in response to pathogen challenges (Ofir et al., 2021; Eastman et al., 2022a; Essuman et al., 2022; Lapin et al., 2022). Deciphering the unique structures and physiological outputs of TIR-signals is progressing, but remains challenging (Wan et al., 2019; Duxbury et al., 2020; Essuman et al., 2022; Huang et al., 2022; Jia et al., 2022; Leavitt et al., 2022). In this report, we generated an auto-active variant of the prokaryotic TIR, ThoerisB (ThsB) and reconstructed Thoeris inputs and outputs *in planta*, to test the hypothesis that plant and prokaryotic signals are conserved. We found that the signals required to activate EDS1 and ThsA were not conserved, and we identify ThsB-generated 3’cADPR as an activator of the Thoeris mediator, ThsA. ThsB-Auto also produces an undefined cADPR-isomer, which remains a potential ThsA-activator. We demonstrate that some plant TIRs simultaneously produce both 2’ and 3’cADPR, and provide a plausible explanation for the cross-kingdom activation of the Thoeris system by the plant BdTIR. Indeed, plant TIRs can generate numerous types of putative signals, and even utilize nucleotide substrates beyond NAD^+^ (Huang et al., 2022; Jia et al., 2022; Manik et al., 2022; Yu et al., 2022).

The detection of bacteriophage triggers ThsB NADase activities, though how ThsB mechanistically transitions to an active state is not understood (Ofir et al., 2021). Plant TIR-NLRs are also activated by pathogen detection, and cryo-EM studies indicate that a conserved ‘BB-loop’ shifts during activation to influence substrate access and TIR-TIR associations (Ma et al., 2020; Martin et al., 2020). This conserved TIR loop is also critical for the catalytic activity of the prokaryotic virulence factor, AbTir (Manik et al., 2022). Modifying this analogous loop within ThsB generates an auto-active variant; however, it is unknown if phage detection effects similar conformational changes within ThsB during activation. Amongst the plant TIR-domains tested, similar BB-loop alterations had the opposite phenotype, resulting in a loss of function. This indicates that plant and prokaryotic TIRs have distinct requirements for catalysis.

During the preparation of this manuscript, two independent studies on the Thoeris system were posted to Biorxiv by Leavitt et al. and Manik et al., respectively (Leavitt et al., 2022; Manik et al., 2022). In contrast to Manik et al. and the report here, Leavitt et al. identifies 2’cADPR as the Thoeris activation signal (Leavitt et al., 2022). Leavitt also identified a phage-encoded repressor of Thoeris, Tad1 (Thoeris anti-defense 1), which blocks ThsA-activation by sequestering cADPR-isomers. Although a Tad1 structure with bound 2’cADPR was generated, this cADPR-signal was derived from BdTIR and not from ThsB. Critically, Tad1’s specificity for different cADPR-isomers has not been resolved, and Tad1 could sequester both 3’cADPR and 2’cADPR (Leavitt et al., 2022). Thus, while we identify 3’cADPR-producing TIRs (ThsB-Auto, ThsB2, HopAM1, BdTIR) to effectively signal to ThsA, our findings do agree that 2’cADPR is the major NADase product of BdTIR (Leavitt et al., 2022). Importantly, we find that BdTIR also generates 3’cADPR in minor amounts, which explains the cross-activation of ThsA reported by Ofir et al. (Ofir et al., 2021). In the report of Manik et al., ThsB products were not directly examined, but rather, purified isomers of 2’ and 3’cADPR were assayed for ThsA stimulation (Manik et al., 2022). Consistent with our studies, an *Enterococcus faecium* encoded ThsA-ortholog (*Ef*ThsA) was found to be ~100-fold more sensitive to 3’cADPR (Manik et al., 2022). It remains possible that ThsA might be further stimulated by the second, and structurally unresolved, cADPR-isomer that we detected from ThsB-Auto *in vitro*.

Similar to *in vitro* reports, BdTIR cross-activated ThsA in our *in planta* assays indicating some conservation of signaling (Ofir et al., 2021). However, in the reciprocal experiment, ThsB-Auto did not cross-activate plant EDS1-dependent TIR-pathways. Recent reports find that the plant TIR-produced metabolites pRib-AMP and ADPr-ATP promote EDS1-family heterodimer formation and association with the downstream helper NLRs, ADR1 and NRG1 (Huang et al., 2022; Jia et al., 2022). While it has not been formally demonstrated that 2’cADPR is dispensable for EDS1 pathway activation *in vivo*, pRib-AMP or ADPRr-ATP appear sufficient *in vitro* to link the EDS1 complex to downstream helper NLR oligomerization (Huang et al., 2022; Jia et al., 2022). Despite this, it remains possible that 2’cADPR and 3’cADPR have alternate functions in plant immunity. Plant TIR-only proteins like BdTIR generate ~100-fold more v-cADPR (2’cADPR) *in planta* relative to TIR-domains from TNL proteins (Wan et al., 2019). In addition, like ThsB-Auto and HopAM1, BdTIR also generates 3’cADPR, although to a lesser extent than 2’cADPR. Given that 3’cADPR is produced by the phytopathogen effector HopAM1, if it is a signaling molecule, it presumably might act as a negative regulator of plant immunity. Curiously, plant TIR-only proteins also generate 2’,3’cNMPs, which can regulate the formation of stress granules (Kosmacz et al., 2018; Yu et al., 2022). Future studies will examine how TIR-generated 3’cADPR or 2’,3’cNMP impact stress or associated transcriptional defense responses, as well as if elevated levels of 2’cADPR could have signaling roles independent of *EDS1*.

In addition to 3’cADPR, we detected an unknown isomer of cADPR that ThsB-Auto generates. The discovery of Tad1 indicates that phages are evolving to circumvent the Thoeris system (Leavitt et al., 2022). Accordingly, Thoeris systems are under selective pressure to overcome Tad1 inhibition. Certain Thoeris operons encode several ThsB paralogs, though it is not clear if ThsB-stacking might further benefit immunity beyond potentially enabling the detection of multiple phage types (Doron et al., 2018). For instance, an expanded ThsB arsenal could enable enzymatic outputs to diversify and produce new signal molecule types and/or have altered kinetics. ThsA orthologs have varying sensitivity to different TIR-derived products (Manik et al., 2022). ThsA-activation by particular cADPR-isomers might also be under selection in response to Tad1-containing phages (Leavitt et al., 2022). It will be interesting to assess ThsA-stimulation by the unknown cADPR-isomer produced by ThsB-Auto, and if Tad1 similarly blocks this putative signal as well.

We do not yet know the mechanism by which BB-loop modification enables ThsB auto-activity. Similar substation of glycine residues within the BB-loop of a prokaryotic TIR-STING protein results in a loss of NADase functions (Morehouse et al., 2022). However, TIR-STING proteins lack apparent cyclase activity, suggesting differences in the catalytic mechanism, as compared to ThsB TIRs (Morehouse et al., 2022). ThsB2-TIR did not require BB-loop modifications to stimulate ThsA, suggesting that ThsB and ThsB2 NADase activities maybe differentially regulated. While BB-loop alteration confers ThsB auto-activity, it is possible this change could shunt NADase outputs from 3’-cADPR-generation towards production of the unknown cADPR-isomer. The finding that HopAM1 and ThsB2 can both stimulate ThsA but do not produce the structurally undefined isomer made by ThsB-Auto, indicates that this isomer is at least not required for ThsA-activation. Regardless, these findings indicate that the targeted alteration of TIR BB-loops might allow for the generation of new and potentially useful products by TIRs.

Unstimulated ThsA has detectable NADase activities *in vitro* (Ka et al., 2020; Ofir et al., 2021; Manik et al., 2022). While our *in planta* reconstruction showed that ThsA by itself does diminish NAD^+^ levels relative to the catalytically inactive SIR2-mutant (N112A), cytotoxic phenotypes were only apparent after ThsA-stimulation by ThsB-Auto. Manik et al. assayed other ThsA-orthologs and found that *Ef*ThsA (*E. faecium*) and *Se*ThsA (*S. equi*) had lower unstimulated NADase activities *in vitro* relative to the prototypic ThsA of *B. cereus* MSX-D12 (Manik et al., 2022). It is possible that this unstimulated ‘background’ NADase activity of ThsA *in vitro* is influenced by recombinant protein preparation. Whether Thoeris systems impose host fitness penalties is not clear (Bernheim and Sorek, 2020); however, because ThsA can be readily expressed in *E. coli*, it presumably is not strongly cytotoxic to host cells (Ka et al., 2020; Ofir et al., 2021; Leavitt et al., 2022; Manik et al., 2022). Further studies could examine whether the SLOG-domain, in absence of ThsB-signals, is influenced by other cellular metabolites or regulators which act to repress unstimulated ThsA NADase activities.

We generated ThsA R371A, which reportedly prevents stimulation by ThsB-generated signals (Ofir et al., 2021; Manik et al., 2022). However, R371A, as well as SLOG substitutions at E403 or K388, resulted in a constitutive enhancement of NADase activity that was cytotoxic *in planta*. A recent crystal structure of the ThsA SLOG-domain shows that these three residues coordinate 3’cADPR binding and subsequently influence NAD^+^ access to the catalytic SIR2-domain (Manik et al., 2022). Within *Ef*ThsA, substituting these residues resulted in a loss of sensitivity to 3’cADPR, at least *in vitro* (Manik et al., 2022). Why mutating these residues within ThsA (*B. cereus* MSX D-12) promotes auto-activity *in planta* is unclear, though it is possible these substitutions could enhance ThsA sensitivity to other metabolites within plant cells (Manik et al., 2022). Regardless, we provide evidence that the SLOG-domain likely has roles in both inhibition and activation. Whether SLOG-domains from phylogenetically distant ThsA proteins are equally stimulated by 3’cADPR vs. other TIR-derived metabolites is not known. Analyzing the conservation of SLOG-residues among ThsA-orthologs may provide clues into whether other Thoeris systems might use unique ThsB-generated signals.

Enzymatic TIRs are prevalent in the immune systems of prokaryotes and eukaryotes, and their roles are highly adaptable. For instance, Thoeris TIRs generate immune signals, while other TIRs act as executioners that simply deplete NAD^+^ (Burroughs et al., 2015; Doron et al., 2018; Hogrel et al., 2022; Koopal et al., 2022; Morehouse et al., 2022). Recently, NLR-like immune proteins were reported in prokaryotes, and certain unicellular algae encode TIR-NB-LRR proteins (Sun et al., 2014; Shao et al., 2019; Kibby et al., 2022). Evolution adapts existing genetic modules into new roles, and whether plant TIR-pathways can be directly traced to particular bacterial or unicellular eukaryote lineages remains an open question (Gao et al., 2022). Further study of the TIR-domains encoded by prokaryotes and early plant lineages will shed light on the conservation of TIR-signals, and likely reveal new types of TIR-generated immune signals.

## Materials & Methods

### Immunoblots

Three 6 mm discs from three different transformed *Nb* leaves were harvested into a 2.0 mL tube containing a single glass bead and flash-frozen in liquid nitrogen. Tissue was homogenized in a TissueLyzer II (Qiagen) at 30 Hz for 30 seconds. Proteins were extracted in 200 μL of lysis buffer [50 mM Tris·HCl (pH 7.5), 150 mM NaCl, 5 mM EDTA, 0.2% Triton X-100, 10% (vol/vol) glycerol, 1/100 Sigma protease inhibitor cocktail]. Lysates were centrifuged at 4°C for 5 minutes at 5k RPM and stored on ice. Equal volumes of lysates were loaded in each sample lane for analysis by SDS/PAGE. After transfer, nitrocellulose blots were blocked for 1 h at RT in 3% (wt/vol) nonfat dry milk TBS-T (50 mM Tris, 150 mM NaCl, 0.05% Tween 20). Immunoblots for either HA (3F10, Roche) or GFP (Cat# 1181446001, Roche) were incubated for 1 h at RT in 3% (wt/vol) nonfat dry milk TBS-T at 1:2,000, followed by 3X washes in TBS-T. Secondary HRP-conjugated goat anti-rabbit or goat anti-mouse IgG (Milipore Sigma) was added at 1:10,000 in TBS-T (3% milk) and incubated for 1 h at room temperature on a platform shaker, followed by 3X washes with TBS-T. Chemiluminescence detection performed with Clarity Western ECL Substrate (Bio-Rad) and developed using a ChemiDoc MP chemiluminescent imager (Bio-Rad).

### Transient Agrobacterium Expression in *Nicotiana benthamiana*

*Agrobacterium tumefaciens* strain GV3101 was syringe infiltrated at OD_600_ of 0.80 into young leaves of ~4-5 wk old *N. benthamiana* plants. Viral suppressor of silencing p19 was included within *Nb* infiltrations at OD 0.05, as described previously (Nishimura et al., 2017; Wan et al., 2019). GV3101 liquid cultures were grown overnight at 28°C in 50 μg/mL rifampicin, gentamicin and spectinomycin. Cultures were induced ~3 h in 10 mM MES buffer (pH 5.60), 10 mM MgCl_2_, and 100 μM acetosyringone prior to infiltration. *N. benthamiana* plants were grown in a Percival set at 25°C with a photoperiod of 16 h light at 80 μE·m−2·s−1. For Thoeris system reconstruction assays, GV3101 cultures at OD 0.80 were mixed 1:1 with respective constructs prior to co-infiltration, unless otherwise noted in Figure panel.

#### Protein Structure Modeling

Protein structure homology models for full-length AbTir and TcpO proteins were generated using AlphaFold (Mirdita et al., 2022) and resulting PDB files were analyzed with PyMOL (The PyMOL Molecular Graphics System, Version 1.8 Schrödinger, LLC). ThsB (PDB ID 6LHY) and ThsA (PDB ID: 6LHX) (*B. cereus* MSX D-12) structures were previously generated by (Ka et al., 2020). RPP1 (PDB ID: 7CRC) cryo-EM structure was generated by (Ma et al., 2020).

#### DNA Vector Construction & PCR Mutagenesis

DNA fragments encoding codon optimized ThsB and ThsA (from *B. cereus* MSX D-12) and other ThsB-orthologs were synthesized by Twist Biosciences with BP-compatible recombination ends. All DNA fragments were BP-cloned into vector pDNR207 using Gateway™ BP Clonase (Invitrogen) and sequence verified (GeneWiz). ORFs were cloned using Gateway™ LR Clonase (Invitrogen) into previously described binary vectors containing Omega leader sequences: pGWB602 (35S Omega:), pGWB615 (35S Omega: HA-), pGWB641 (35S Omega: -GFP) or pGWB602 (35S: HA_SAM-). The polymerase incomplete primer extension (PIPE) method was used to introduce single amino acid substitutions or small insertions/deletions (Klock and Lesley, 2009). PCR was performed on genes within pDNR207 constructs with Q5 High-Fidelity polymerase (New England Biolabs, Ipswich MA), and all constructs were sequence verified by GeneWiz prior to cloning into binary vectors. The HA_SAM oligomerization domain (1X HA tag-SARM1_478-578_-GGGGS) of SARM1, the SARM1-TIR domain (561-724), and the TcpO-TIR domain (residues 204-341) were described previously by Wan et al., 2019. The AbTir core-TIR domain (AbTIR) corresponds to residues 134-269. Amino acid sequences for TIR and ThsA constructs used is provided in the Supporting Information.

#### Recombinant Protein Expression and Purification

Corresponding ThsB, AbTIR, HopAM1 and BdTIR constructs were cloned into the pET30a+ vector with N-terminal Strep II tag and C-terminal 6XHis tag. Proteins were expressed in E. coli BL21(DE3) and induced overnight using the auto-induction method (Studier, 2005). Cell pellets were resuspended in lysis buffer (50 mM HEPES pH 8.0, 300 mM NaCl and 1 mM DTT) and lysed using sonication. The resulting supernatant was filtered and applied onto NiA-beads. The beads were washed with lysis buffer. Bound proteins were eluted using elution buffer (50 mM HEPES pH 8.0, 300 mM NaCl and 250 mM imidazole). Eluted proteins were concentrated and applied onto a Superdex 75 HiLoad 26/60 size-exclusion column (GE Healthcare) pre-equilibrated with gel-filtration buffer (10 mM HEPES pH 7.5, 150 mM NaCl, 1 mM DTT). The peak fractions were pooled and confirmed by SDS-PAGE and concentrated using Amicon Ultra-15 Centrifugal Filter Units (Millipore). The protein samples were stored at −80°C.

#### *In vitro* NADase Assays

Ten microliters of NiA-beads bound with purified proteins were incubated with 30 μM NAD^+^ (final concentration) in 50 μL buffer (92.4 mM NaCl and 0.64X PBS). Reactions were carried out at room temperature (around 25° C) for 2 h and stopped by addition of 50 μL of 1M perchloric acid (HClO_4_) and then placed on ice for 10 min. Neutralization was performed by adding 16.7 μL of 3M K_2_CO_3_. Samples were placed on ice for another 10 min, and then cleared by centrifugation. Extracted metabolites were stored at −20°C until later LC-MS/MS analysis.

#### *In vitro* LC-MS/MS Metabolite Measurements

The samples were centrifuged at 12,000 × g for 10 min, and the cleared supernatant was applied to the LC-MS/MS for metabolite identification and quantification. Samples were analyzed by Q Exactive quadrupole orbitrap high resolution mass spectrometry coupled with a Dionex Ultimate 3000 RSLC (HPG) ultra-performance liquid chromatography (UPLC-Q-Orbitrap-HRMS) system (Thermo Fisher Scientific) and a HESI ionization source under positive ion modes using a heated ESI source. The injection volume was set to 1 μL. Samples were separated with an ACQUITY UPLC HSS T3 column (100 mm × 2.1 mm, 1.8-μm particle size; Waters). The mobile phase consisted of 2 mM ammonium formate in water (A) and in 100% methanol (B), and the gradient elution was set as following: 0–2.00 min, 1% B; 2.00–7.00 min, 1%–95% B; 9.00–9.10 min, 95%–1% B; 9.10–11.00 min, 1% B. The flow rate was set to 0.3 mL/min and column temperature at 45°C. Metabolites were quantified by using area and the retention time for each compound was determined using standards including NAD^+^, cADPR and Nam dissolved in 50% methanol. The metabolites were also detected and quantified with a Triple Quad mass spectrometer (6500; Agilent) under positive ESI multiple reaction monitoring (MRM) for monitoring analyte parent ion and product ion formation. MRM conditions were optimized using authentic standard chemicals including: NAD^+^ ([M+H]+ 664 > 136.00, 664 > 428, 664 > 542); Nam ([M+H]+ 123 > 80); cADPR ([M+H]+ 542 > 136, 542 > 348, 542 >428); v-cADPR ([M+H]+ 542 >136).

#### *In planta* LC-MS Analysis

LCMS/MS analysis was carried out on Waters TQ-XS triple quad mass spectrometer coupled with Waters H-class UPLC. Column used is a Waters Acquity UPLC-C18, 1.7 μm, 2.1×50mm. Mobile phases include A: 2mM ammonium acetate and B: methanol. Flow rate: 0.2 mL/min. Gradient: 0-5 min, 0% B; 5-7min, 0% to 20% B; 7-8min, 20% to 100% B; then the column is washed with 100% B for 2 mins before equilibration to 100% A for 15 mins. Mass spectrometer conditions: Capillary voltage: 800V; desolvation temperature: 600°C; desolvation gas (nitrogen, 1000 L/hr); cone gas: 150 L/hr, Nebuliser gas: 7 bar. MRM parameters for the detection of cADPR-isomers, 542/136 (cone voltage 20V, collision energy: 32eV); 542/348 (cone voltage 20V, collision energy: 28eV). cADPR-isomers were verified by comparing with authentic standards, including 2’cADPR and 3’cADPR.

#### *In planta* NADase Assays

Three 6 mm discs of transformed *Nb* leaf tissue were harvested into 2.0 mL tubes containing a single glass bead and flash-frozen in liquid nitrogen. Tissue was homogenized in a Qiagen TissueLyzer II at 30 Hz for 30 seconds. Tissue was resuspended in 300 μL of ice-cold lysis buffer [50 mM Tris·HCl (pH 7.5), 150 mM NaCl, 5 mM EDTA, 0.2% Triton X-100 and 10% (vol/vol) glycerol] and stored on ice. Lysates were centrifuged at 4°C for 5 minutes at 5k RPM and returned on ice. Supernatant was diluted 1:3 in lysis buffer before addition to Amplite NAD^+^-Assay kit (Cat# 15280, AAT Biosciences). Colorimetric reagent was developed for 20 minutes at room temperature. Colorimetric NAD^+^-detection was performed in 96-well black bottom plates (Costar) on an Infinite M Plex (Tecan) plate reader at excitation (420 nm) and emission (480 nm) using i-control 2.0 software.

#### Ion Leakage Assays

Ion leakage assays were performed essentially as described in (Nishimura et al., 2017). Briefly, 6x 4 mm leaf discs were collected from *Nb* plants at ~3 days after *Agro*-infiltration and placed into sterile 15-mL conical tubes containing 6.0 mL of ultrapure H_2_O. Ion measurements were recorded at noted times using an OrionStar A112 conductivity meter (Thermo Scientific).

#### Statistical Analyses

Multiple comparisons were analyzed via one-way ANOVA with post hoc Tukey HSD (honestly significant differences) using R Studio (version 1.4.1717, RStudio Team (2021), Boston, MA). Comparisons of significance indicated with compact letter display (CLD). Over-lapping letters are non-significant (*p*>.05) while separate letter classes indicate *p*<.05 or better.

## Acknowledgements and Funding

We would like to thank Dr. Qingli Liu, Dr. Samuel Eastman, Dr. Farid El-Kasmi, and Dr. Jeffrey Dangl for helpful guidance and suggestions. We also thank Eliza Walthers, Tyler Todd, and Christopher Gomez for assistance in growing and caring for plants and other resources. This work was supported by start-up funds provided by Colorado State University and a National Science Foundation (IOS-1758400) grant awarded to M.T.N. Post-doctoral fellowship support for A.M.B. was provided by Syngenta Crop Protection. B.K. and T.V. were supported by the National Health and Medical Research Council (NHMRC grants 1196590, 1107804, 1160570 1071659 and 1108859), the Australian Research Council (ARC) Future Fellowship (FT200100572) to T.V., the ARC Laureate Fellowship (FL180100109) to B.K., and the ARC DECRA (DE170100783) to T.V. M.G. and L.S. were supported by a grant from the Biotechnology and Biological Sciences Research Council (BB/V00400X/1). L.W. was supported by National Key Laboratory of Plant Molecular Genetics, the Institute of Plant Physiology and Ecology/Center for Excellence in Molecular Plant Sciences, and the Chinese Academy of Sciences Strategic Priority Research Program (type B; project number XDB27040214).

## References

Bayless, A.M., and Nishimura, M.T. (2020). Enzymatic functions for toll/interleukin-1 receptor domain proteins in the plant immune system. Front Genet 11, 539.

Bernheim, A., and Sorek, R. (2020). The pan-immune system of bacteria: antiviral defence as a community resource. Nat Rev Microbiol 18, 113–119.

Bi, G., Su, M., Li, N., Liang, Y., Dang, S., Xu, J., Hu, M., Wang, J., Zou, M., Deng, Y., Li, Q., Huang, S., Li, J., Chai, J., He, K., Chen, Y.H., and Zhou, J.M. (2021). The ZAR1 resistosome is a calcium-permeable channel triggering plant immune signaling. Cell 184, 3528–3541 e3512.

Burroughs, A.M., Zhang, D., Schaffer, D.E., Iyer, L.M., and Aravind, L. (2015). Comparative genomic analyses reveal a vast, novel network of nucleotide-centric systems in biological conflicts, immunity and signaling. Nucleic Acids Res 43, 10633–10654.

Coronas-Serna, J.M., Louche, A., Rodriguez-Escudero, M., Roussin, M., Imbert, P.R.C., Rodriguez-Escudero, I., Terradot, L., Molina, M., Gorvel, J.P., Cid, V.J., and Salcedo, S.P. (2020). The TIR-domain containing effectors BtpA and BtpB from Brucella abortus impact NAD metabolism. PLoS Pathog 16, e1007979.

Doron, S., Melamed, S., Ofir, G., Leavitt, A., Lopatina, A., Keren, M., Amitai, G., and Sorek, R. (2018). Systematic discovery of antiphage defense systems in the microbial pangenome. Science 359(6379).

Duxbury, Z., Wang, S., MacKenzie, C.I., Tenthorey, J.L., Zhang, X., Huh, S.U., Hu, L., Hill, L., Ngou, P.M., Ding, P., Chen, J., Ma, Y., Guo, H., Castel, B., Moschou, P.N., Bernoux, M., Dodds, P.N., Vance, R.E., and Jones, J.D.G. (2020). Induced proximity of a TIR signaling domain on a plant-mammalian NLR chimera activates defense in plants. Proc Natl Acad Sci U S A 117, 18832–18839.

Eastman, S., Bayless, A., and Guo, M. (2022a). The nucleotide revolution: immunity at the intersection of TIR-domains, nucleotides, and Ca^2+^. Mol Plant Microbe Interact.

Eastman, S., Smith, T., Zaydman, M.A., Kim, P., Martinez, S., Damaraju, N., DiAntonio, A., Milbrandt, J., Clemente, T.E., Alfano, J.R., and Guo, M. (2022b). A phytobacterial TIR domain effector manipulates NAD(+) to promote virulence. New Phytol 233, 890–904.

El Kasmi, F. (2021). How activated NLRs induce anti-microbial defenses in plants. Biochem Soc Trans 49, 2177–2188.

Essuman, K., Milbrandt, J., Dangl, J.L., and Nishimura, M.T. (2022). Shared TIR enzymatic functions regulate cell death and immunity across the tree of life. Science 377, eabo0001.

Essuman, K., Summers, D.W., Sasaki, Y., Mao, X., DiAntonio, A., and Milbrandt, J. (2017). The SARM1 Toll/interleukin-1 receptor domain possesses intrinsic NAD(+) cleavage activity that promotes pathological axonal degeneration. Neuron 93, 1334–1343 e1335.

Essuman, K., Summers, D.W., Sasaki, Y., Mao, X., Yim, A.K.Y., DiAntonio, A., and Milbrandt, J. (2018). TIR domain proteins are an ancient family of NAD(+)-consuming enzymes. Curr Biol 28, 421–430 e424.

Ficke, A., Cowger, C., Bergstrom, G., and Brodal, G. (2018). Understanding Yield Loss and Pathogen Biology to Improve Disease Management: Septoria Nodorum Blotch - A Case Study in Wheat. Plant disease 102, 696–707.

Fliegert, R., Riekehr, W.M., and Guse, A.H. (2020). Does Cyclic ADP-Ribose (cADPR) Activate the Non-selective Cation Channel TRPM2? Front Immunol 11, 2018.

Gallie, D.R. (2002). The 5’-leader of tobacco mosaic virus promotes translation through enhanced recruitment of eIF4F. Nucleic Acids Res 30, 3401–3411.

Gao, L.A., Wilkinson, M.E., Strecker, J., Makarova, K.S., Macrae, R.K., Koonin, E.V., and Zhang, F. (2022). Prokaryotic innate immunity through pattern recognition of conserved viral proteins. Science 377, eabm4096.

Gerdts, J., Brace, E.J., Sasaki, Y., DiAntonio, A., and Milbrandt, J. (2015). SARM1 activation triggers axon degeneration locally via NAD(+) destruction. Science 348, 453–457.

Hogrel, G., Guild, A., Graham, S., Rickman, H., Gruschow, S., Bertrand, Q., Spagnolo, L., and White, M.F. (2022). Cyclic nucleotide-induced helical structure activates a TIR immune effector. Nature 608(7924), 808–812.

Horsefield, S., Burdett, H., Zhang, X., Manik, M.K., Shi, Y., Chen, J., Qi, T., Gilley, J., Lai, J.S., Rank, M.X., Casey, L.W., Gu, W., Ericsson, D.J., Foley, G., Hughes, R.O., Bosanac, T., von Itzstein, M., Rathjen, J.P., Nanson, J.D., Boden, M., Dry, I.B., Williams, S.J., Staskawicz, B.J., Coleman, M.P., Ve, T., Dodds, P.N., and Kobe, B. (2019). NAD(+) cleavage activity by animal and plant TIR domains in cell death pathways. Science 365, 793–799.

Huang, S., Jia, A., Song, W., Hessler, G., Meng, Y., Sun, Y., Xu, L., Laessle, H., Jirschitzka, J., Ma, S., Xiao, Y., Yu, D., Hou, J., Liu, R., Sun, H., Liu, X., Han, Z., Chang, J., Parker, J.E., and Chai, J. (2022). Identification and receptor mechanism of TIR-catalyzed small molecules in plant immunity. Science 377, eabq3297.

Jacob, P., Kim, N.H., Wu, F., El-Kasmi, F., Chi, Y., Walton, W.G., Furzer, O.J., Lietzan, A.D., Sunil, S., Kempthorn, K., Redinbo, M.R., Pei, Z.M., Wan, L., and Dangl, J.L. (2021). Plant “helper” immune receptors are Ca(2+)-permeable nonselective cation channels. Science 373, 420–425.

Jia, A., Huang, S., Song, W., Wang, J., Meng, Y., Sun, Y., Xu, L., Laessle, H., Jirschitzka, J., Hou, J., Zhang, T., Yu, W., Hessler, G., Li, E., Ma, S., Yu, D., Gebauer, J., Baumann, U., Liu, X., Han, Z., Chang, J., Parker, J.E., and Chai, J. (2022). TIR-catalyzed ADP-ribosylation reactions produce signaling molecules for plant immunity. Science 377, eabq8180.

Johanndrees, O., Baggs, E.L., Uhlmann, C., Locci, F., Läßle, H.L., Melkonian, K., Käufer, K., Dongus, J.A., Nakagami, H., Krasileva, K.V., Parker, J.E., and Lapin, D. (2021). Differential *EDS1* requirement for cell death activities of plant TIR-domain proteins. bioRxiv, 2021.2011.2029.470438.

Jones, J.D., Vance, R.E., and Dangl, J.L. (2016). Intracellular innate immune surveillance devices in plants and animals. Science 354(6316).

Ka, D., Oh, H., Park, E., Kim, J.H., and Bae, E. (2020). Structural and functional evidence of bacterial antiphage protection by Thoeris defense system via NAD(+) degradation. Nat Commun 11, 2816.

Kibby, E.M., Conte, A.N., Burroughs, A.M., Nagy, T.A., Vargas, J.A., Aravind, L., and Whiteley, A.T. (2022). Bacterial NLR-related proteins protect against phage. bioRxiv, 2022.2007.2019.500537.

Kim, N.H., Jacob, P., and Dangl, J.L. (2022). Con-Ca(2+) -tenating plant immune responses via calcium-permeable cation channels. New Phytol 234, 813–818.

Klock, H.E., and Lesley, S.A. (2009). The Polymerase Incomplete Primer Extension (PIPE) method applied to high-throughput cloning and site-directed mutagenesis. Methods Mol Biol 498, 91–103.

Koopal, B., Potocnik, A., Mutte, S.K., Aparicio-Maldonado, C., Lindhoud, S., Vervoort, J.J.M., Brouns, S.J.J., and Swarts, D.C. (2022). Short prokaryotic Argonaute systems trigger cell death upon detection of invading DNA. Cell 185, 1471–1486 e1419.

Kosmacz, M., Luzarowski, M., Kerber, O., Leniak, E., Gutierrez-Beltran, E., Moreno, J.C., Gorka, M., Szlachetko, J., Veyel, D., Graf, A., and Skirycz, A. (2018). Interaction of 2’,3’-cAMP with Rbp47b Plays a Role in Stress Granule Formation. Plant Physiol 177, 411–421.

Lapin, D., Johanndrees, O., Wu, Z., Li, X., and Parker, J.E. (2022). Molecular innovations in plant TIR-based immunity signaling. Plant Cell 34, 1479–1496.

Leavitt, A., Yirmiya, E., Amitai, G., Lu, A., Garb, J., Morehouse, B.R., Hobbs, S.J., Kranzusch, P.J., and Sorek, R. (2022). Viruses inhibit TIR gcADPR signaling to overcome bacterial defense. bioRxiv, 2022.2005.2003.490397.

Li, Y., Pazyra-Murphy, M.F., Avizonis, D., de Sa Tavares Russo, M., Tang, S., Chen, C.Y., Hsueh, Y.P., Bergholz, J.S., Jiang, T., Zhao, J.J., Zhu, J., Ko, K.W., Milbrandt, J., DiAntonio, A., and Segal, R.A. (2022). Sarm1 activation produces cADPR to increase intra-axonal Ca++ and promote axon degeneration in PIPN. J Cell Biol 221(2): e202106080.

Ma, S., Lapin, D., Liu, L., Sun, Y., Song, W., Zhang, X., Logemann, E., Yu, D., Wang, J., Jirschitzka, J., Han, Z., Schulze-Lefert, P., Parker, J.E., and Chai, J. (2020). Direct pathogen-induced assembly of an NLR immune receptor complex to form a holoenzyme. Science 370(6521): eabe3069.

Manik, M.K., Shi, Y., Li, S., Zaydman, M.A., Damaraju, N., Eastman, S., Smith, T.G., Gu, W., Masic, V., Mosaiab, T., Weagley, J.S., Hancock, S.J., Vasquez, E., Hartley-Tassell, L., Kargios, N., Maruta, N., Lim, B.Y.J., Burdett, H., Landsberg, M.J., Schembri, M.A., Prokes, I., Song, L., Grant, M., DiAntonio, A., Nanson, J.D., Guo, M., Milbrandt, J., Ve, T., and Kobe, B. (2022). Cyclic ADP ribose isomers: Production, chemical structures, and immune signaling. Science, eadc8969.

Martin, R., Qi, T., Zhang, H., Liu, F., King, M., Toth, C., Nogales, E., and Staskawicz, B.J. (2020). Structure of the activated ROQ1 resistosome directly recognizing the pathogen effector XopQ. Science 370(6521): eabd9993.

Mirdita, M., Schütze, K., Moriwaki, Y., Heo, L., Ovchinnikov, S., and Steinegger, M. (2022). ColabFold: making protein folding accessible to all. Nature Methods 19, 679–682.

Morehouse, B.R., Yip, M.C.J., Keszei, A.F.A., McNamara-Bordewick, N.K., Shao, S., and Kranzusch, P.J. (2022). Cryo-EM structure of an active bacterial TIR-STING filament complex. Nature 608(7924): 803–807.

Nandety, R.S., Caplan, J.L., Cavanaugh, K., Perroud, B., Wroblewski, T., Michelmore, R.W., and Meyers, B.C. (2013). The role of TIR-NBS and TIR-X proteins in plant basal defense responses. Plant Physiol 162, 1459–1472.

Nimma, S., Ve, T., Williams, S.J., and Kobe, B. (2017). Towards the structure of the TIR-domain signalosome. Curr Opin Struct Biol 43, 122–130.

Nishimura, M.T., Anderson, R.G., Cherkis, K.A., Law, T.F., Liu, Q.L., Machius, M., Nimchuk, Z.L., Yang, L., Chung, E.H., El Kasmi, F., Hyunh, M., Osborne Nishimura, E., Sondek, J.E., and Dangl, J.L. (2017). TIR-only protein RBA1 recognizes a pathogen effector to regulate cell death in Arabidopsis. Proc Natl Acad Sci U S A 114, E2053–E2062.

Ofir, G., Herbst, E., Baroz, M., Cohen, D., Millman, A., Doron, S., Tal, N., Malheiro, D.B.A., Malitsky, S., Amitai, G., and Sorek, R. (2021). Antiviral activity of bacterial TIR domains via immune signalling molecules. Nature 600, 116–120.

Savary, S., Ficke, A., Aubertot, J.-N., and Hollier, C. (2012). Crop losses due to diseases and their implications for global food production losses and food security. Food Security 4, 519–537.

Shao, Z.Q., Xue, J.Y., Wang, Q., Wang, B., and Chen, J.Q. (2019). Revisiting the Origin of Plant NBS-LRR Genes. Trends Plant Sci 24, 9–12.

Shi, Y., Kerry, P.S., Nanson, J.D., Bosanac, T., Sasaki, Y., Krauss, R., Saikot, F.K., Adams, S.E., Mosaiab, T., Masic, V., Mao, X., Rose, F., Vasquez, E., Furrer, M., Cunnea, K., Brearley, A., Gu, W., Luo, Z., Brillault, L., Landsberg, M.J., DiAntonio, A., Kobe, B., Milbrandt, J., Hughes, R.O., and Ve, T. (2022). Structural basis of SARM1 activation, substrate recognition, and inhibition by small molecules. Mol Cell 82, 1643–1659 e1610.

Studier, F.W. (2005). Protein production by auto-induction in high density shaking cultures. Protein Expr Purif 41, 207–234.

Sun, X., Pang, H., Li, M., Chen, J., and Hang, Y. (2014). Tracing the origin and evolution of plant TIR-encoding genes. Gene 546, 408–416.

Tian, L., and Li, X. (2022). TIR domains as two-tiered enzymes to activate plant immunity. Cell 185, 2208–2209.

Wan, L., Essuman, K., Anderson, R.G., Sasaki, Y., Monteiro, F., Chung, E.H., Osborne Nishimura, E., DiAntonio, A., Milbrandt, J., Dangl, J.L., and Nishimura, M.T. (2019). TIR domains of plant immune receptors are NAD(+)-cleaving enzymes that promote cell death. Science 365, 799–803.

Wang, J., Hu, M., Wang, J., Qi, J., Han, Z., Wang, G., Qi, Y., Wang, H.W., Zhou, J.M., and Chai, J. (2019a). Reconstitution and structure of a plant NLR resistosome conferring immunity. Science 364(6435): eaav5870.

Wang, J., Wang, J., Hu, M., Wu, S., Qi, J., Wang, G., Han, Z., Qi, Y., Gao, N., Wang, H.W., Zhou, J.M., and Chai, J. (2019b). Ligand-triggered allosteric ADP release primes a plant NLR complex. Science 364(6435):eaav5868.

Wang, Y., Pruitt, R.N., Nürnberger, T., and Wang, Y. (2022). Evasion of plant immunity by microbial pathogens. Nat Rev Microbiol 20(8), 449–464.

Weagley, J.S., Zaydman, M., Venkatesh, S., Sasaki, Y., Damaraju, N., Yenkin, A., Buchser, W., Rodionov, D.A., Osterman, A., Ahmed, T., Barratt, M.J., DiAntonio, A., Milbrandt, J., and Gordon, J.I. (2022). Products of gut microbial Toll/interleukin-1 receptor domain NADase activities in gnotobiotic mice and Bangladeshi children with malnutrition. Cell Rep 39, 110738.

Yu, D., Song, W., Tan, E.Y.J., Liu, L., Cao, Y., Jirschitzka, J., Li, E., Logemann, E., Xu, C., Huang, S., Jia, A., Chang, X., Han, Z., Wu, B., Schulze-Lefert, P., and Chai, J. (2022). TIR domains of plant immune receptors are 2’,3’-cAMP/cGMP synthetases mediating cell death. Cell 185, 2370–2386 e2318.

