## Supplementary Figures for "Plant and prokaryotic TIR domains generate distinct cyclic ADPR NADase products"

# SI 1

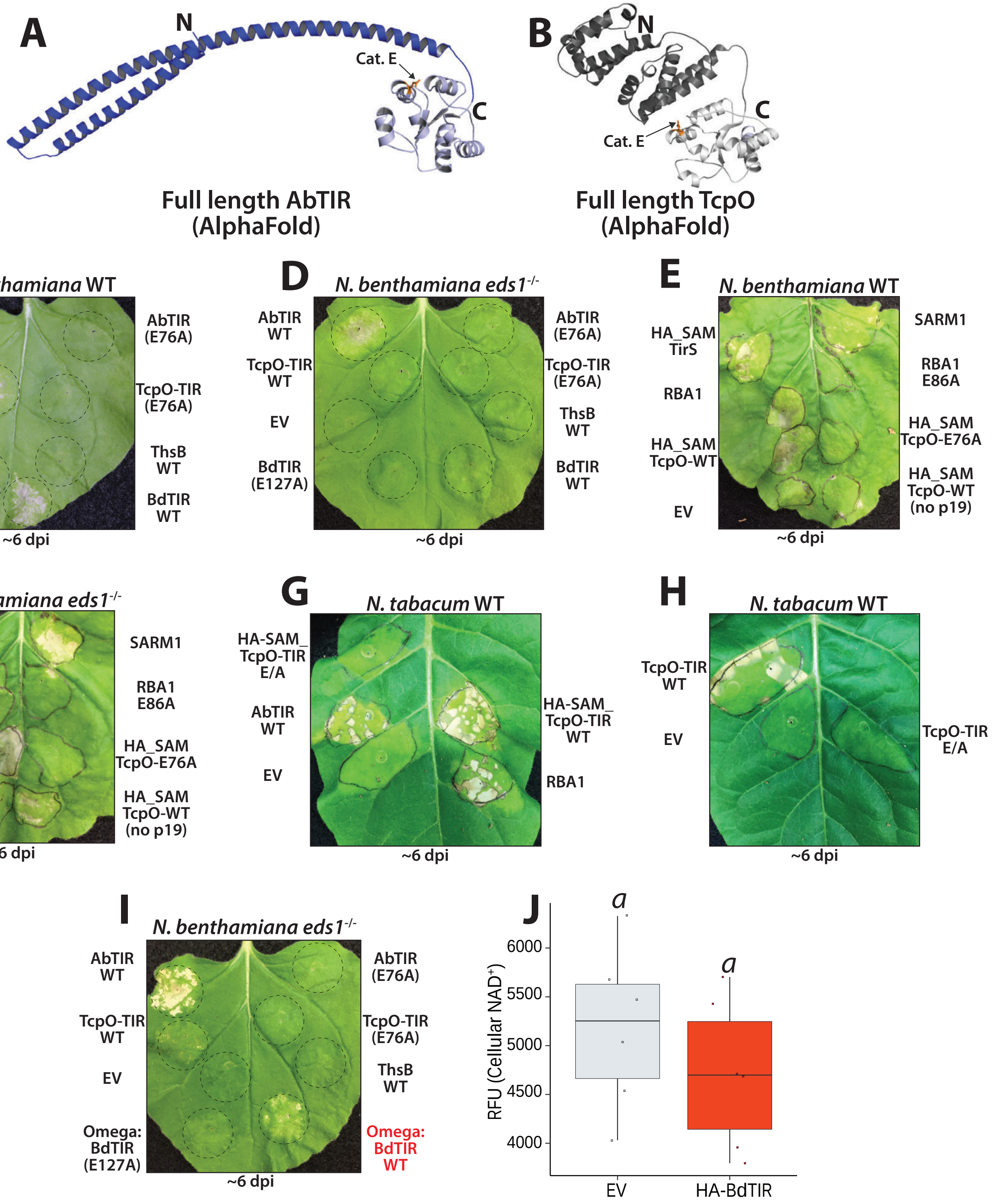


**SI 1**. Over-expression of prokaryotic TIRs can trigger *eds1*$^{-/-}$ independent cell death. (*A* and *B*) AlphaFold models of full length AbTIR encoded by human pathogen, *Acinetobacter baumanii*, or full length TcpO-TIR encoded by *Methanobrevibacter olleyae*. TIR-domain core is colored grey; catalytic glutamate residue (E) is colored orange. (*C-F*) *Nb* WT or *eds1*$^{-/-}$ leaves expressing untagged AbTIR or TcpO-TIR, or HA-SAM-tagged versions of AbTIR, TcpO-TIR or TirS. TirS is a previously described ADPR-producing TIR from *Staphylococcus aureus* (Essuman *et al.*, 2018). SAM (sterile alpha motif of SARM1) oligomerization domain fusions promote TIR-activity, as previously described in Wan *et al*, 2019. EV: 35S-GFP. Positive HR-control RBA1 (Response to HopBA1) has been previously described in Nishimura *et al,* 2017. All constructs infiltrated at OD 0.80 and imaged ~6 dpi. (*G-H*) Like *C*, but *Agro*-infiltration of TIRs into WT *Nicotiana tabacum*. (*I*) 35S Omega leader driven expression of BdTIR can also drive *EDS1*-independent cell death in *Nb*. All constructs infiltrated at OD 0.80 and imaged ~6 dpi. (*J*) Fluorescent NAD$^+$-detection assay in *Nb eds1*$^{-/-}$ leaves performed at 40 hpi. 35S binary constructs expressing 35S:HA-BdTIR or EV (empty vector, 35S: GFP).

# SI 2

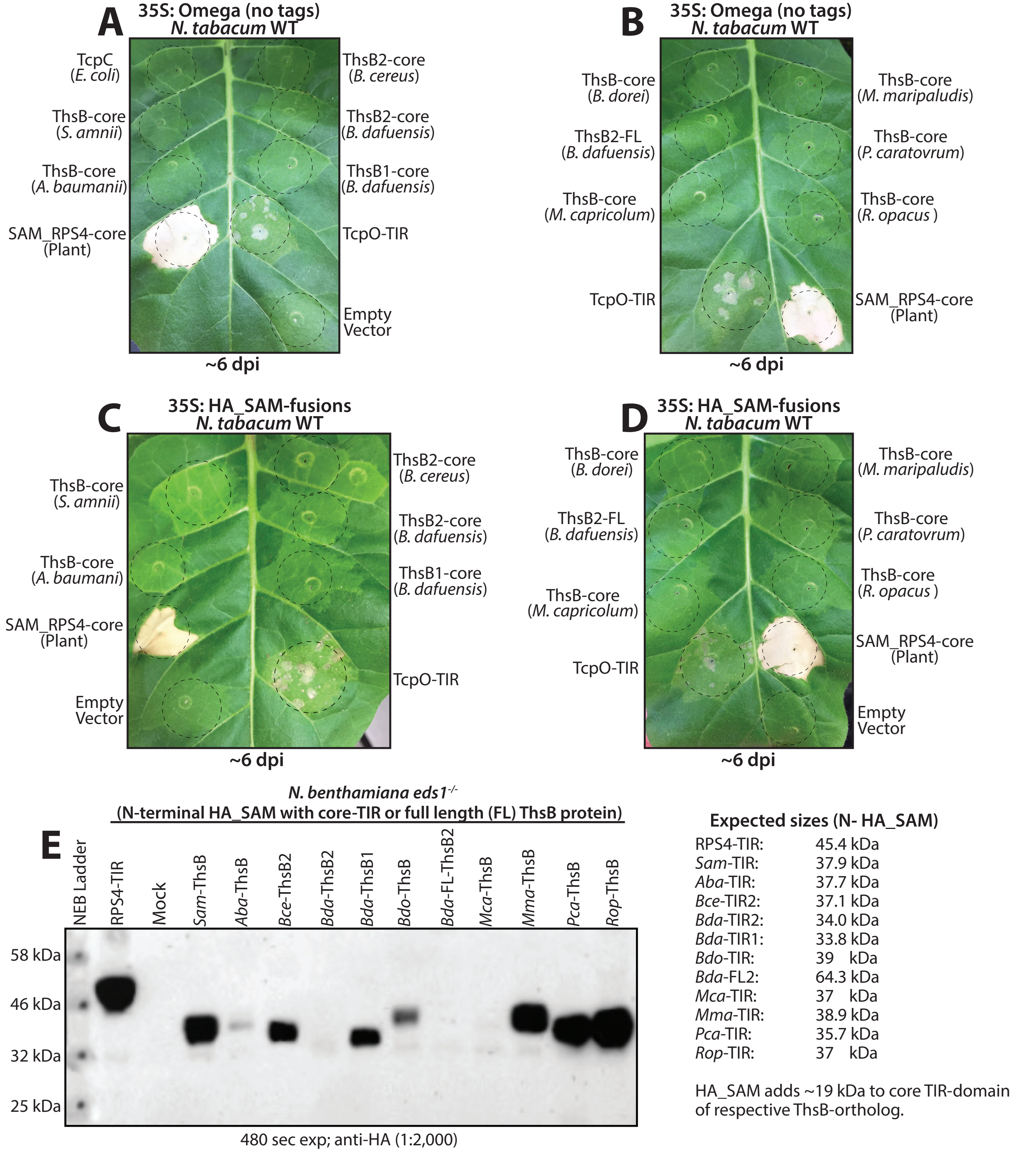


**SI 2**. Examined orthologs of ThsB do not trigger HR. (*A-D*) *Nicotiana tabacum* leaves expressing core TIR-domains (or full-length protein (FL)) of noted ThsB-orthologs, either with or without N-terminal SAM-oligomerization domain fusions. Leaves shown ~5-6 dpi; all constructs infiltrated at OD 0.80. (*E*) Anti-HA immunoblot detection of HA-SAM_ThsB-orthologs harvested from *Nb eds1*$^{-/-}$ leaves at ~40 hpi. Expected size of HA_SAM-fusion proteins and origin of TIR-domain listed on right. Positive HR-control SAM_RPS4-TIR previously described by Wan *et al*., 2019.

# SI3

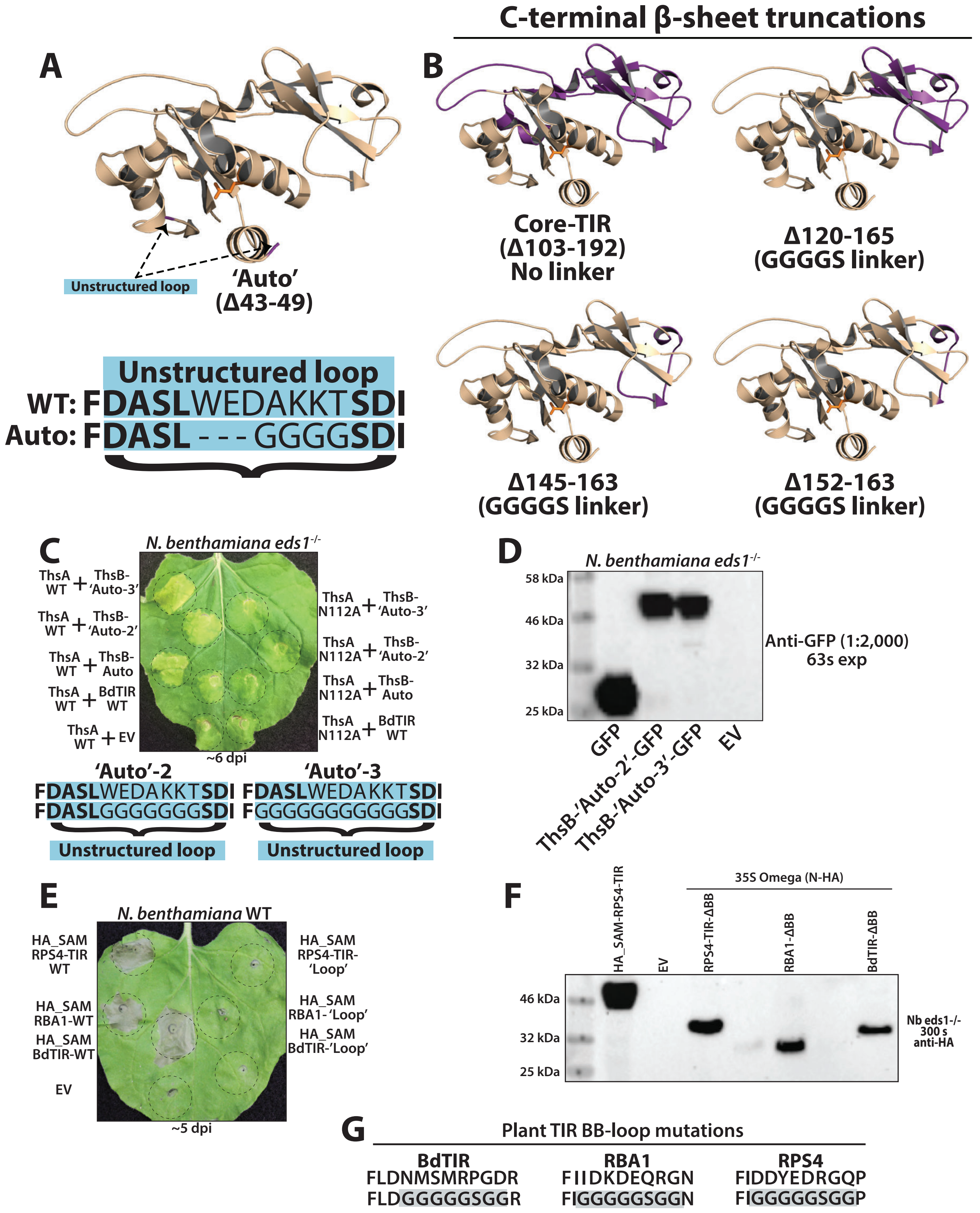


**SI 3**. BB-loop substitutions promote ThsB auto-activity but impair constitutively active plant TIRs. (*A*) Crystal structure of ThsB (PDB ID: 6LHY) determined by Ka *et al*, showing unstructured BB-loop region replaced in the ThsB-Auto active mutant (see arrows) (Ka et al., 2020). Sequence of ThsB-Auto shown below. (*B*) Like A, but structures indicate ThsB C-terminal replacement variants. Replaced C-terminal regions are shown purple. (*C*) *Nb eds1*$^{-/-}$ leaves expressing additional auto-active mutants of ThsB. All constructs infiltrated at OD 0.80 and imaged ~6 dpi. Sequences of mutants shown below. (*D*) Immunoblot of N-HA tagged versions of ThsB-Auto-2 and -3, respectively. (*E*) *Nb* WT leaf indicating phenotypes of plant TIR BB-loop replacement constructs, relative to WT versions. (*F*) Immunoblot of HA-tagged BB-loop placements within plant TIRs. (*G*) Sequence of WT (Top) and BB-loop (bottom) replacements of BdTIR, RBA1 and RPS4.

# SI4

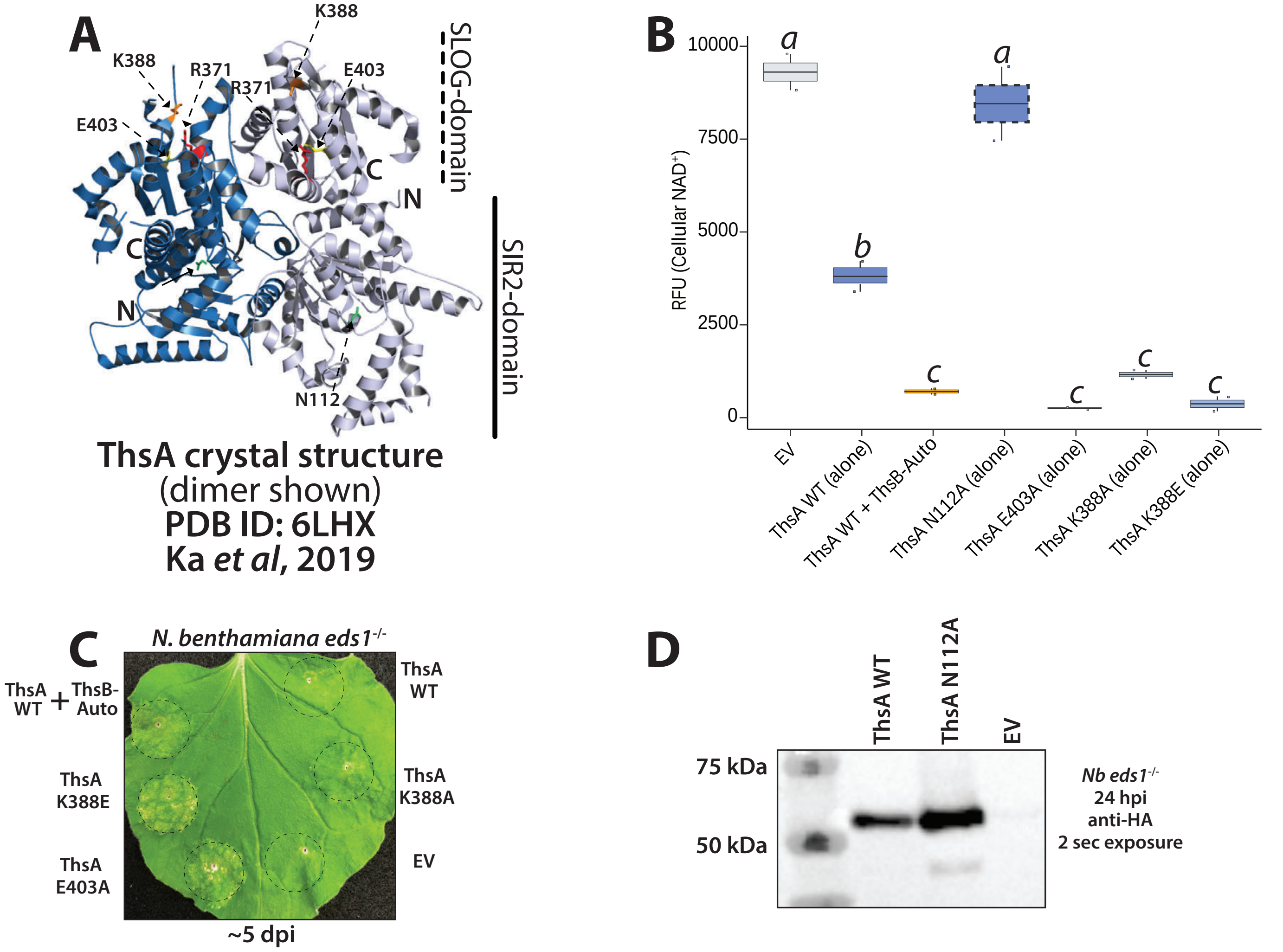


**SI 4**. Substitution of ThsA-SLOG domain residues causes NADase auto-activity and cytotoxicity *in planta* (*A*) Crystal structure of ThsA (PDB ID: 6LHX) determined by Ka *et al* (Ka et al., 2020); two ThsA monomers are shown in blue and grey. N: N-terminus, C: C-terminus. N112 of SIR2 domain shown green, while R371, K388, and E403 within the SLOG-domain are colored red, green or yellow, respectively. (*B*) Fluorescent $NAD^+$-detection assay in *Nb eds1*$^{-/-}$ leaves expressing various ThsA constructs alone, or ThsA + ThsB-Auto. Individual constructs expressed at OD 0.80, and tissue was harvested ~40 hpi. (*C*) *Nb eds1*$^{-/-}$ leaves expressing various auto-active ThsA SLOG-domain constructs. Leaf shown ~5 dpi, and all constructs infiltrated at OD 0.80. (*D*) Anti-HA immunoblot of N-HA tagged ThsA WT or ThsA N112A.

# SI 5

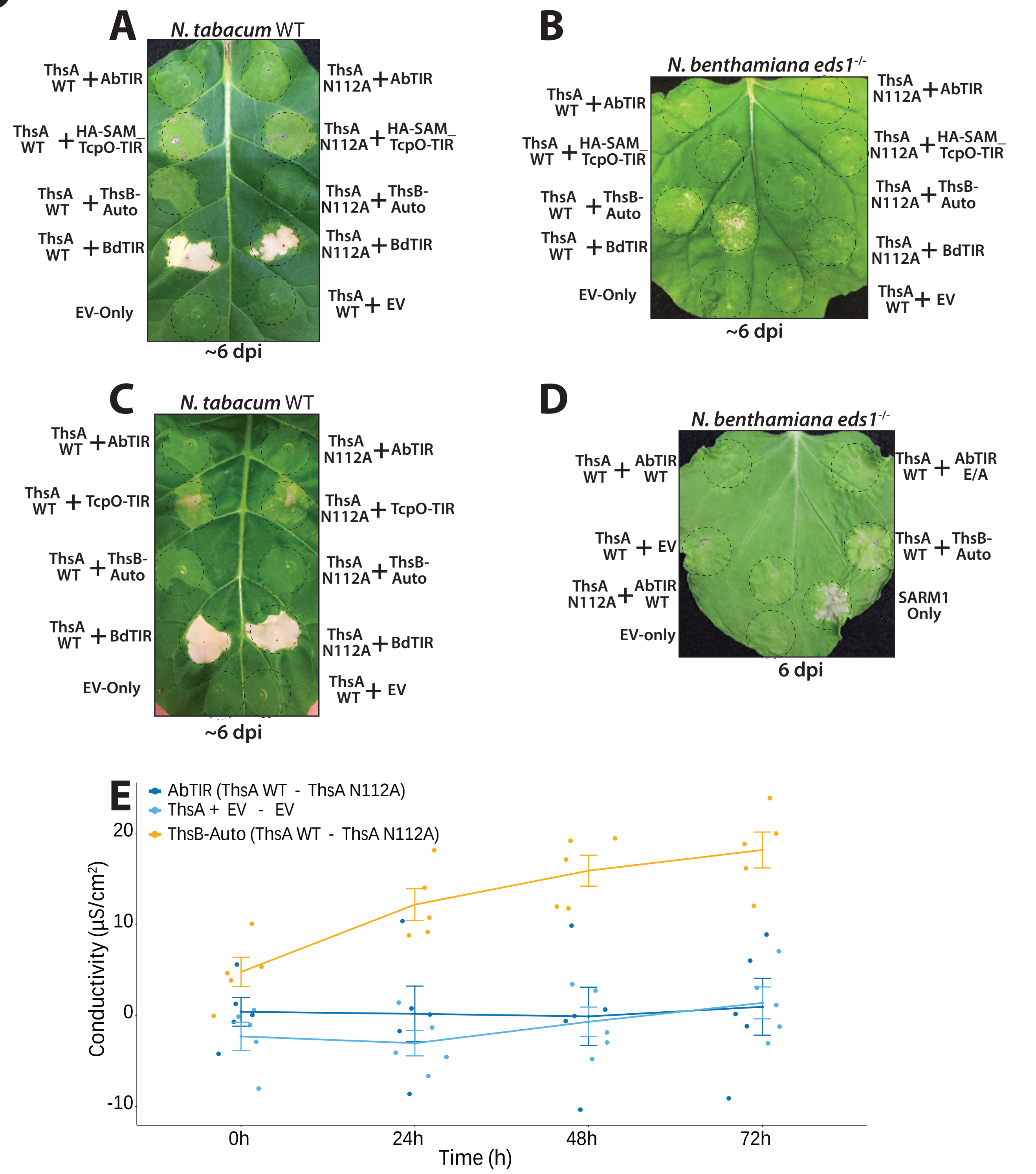


**SI 5**. Co-expression of AbTIR or TcpO-TIR does not enhance stimulation of ThsA activity relative to ThsB-Auto. (*A-D*) *N. tabacum* or *N. benthamiana eds1*$^{-/-}$ leaves co-expressing WT ThsA or N112A in combination with ThsB-Auto, or AbTIR or TcpO-TIR or EV (35S:GFP, empty vector). The SAM-domain promotes oligomerization and refers to the sterile alpha motif of SARM1. ThsA and ThsB variants, or SARM1 or EV (35S:GFP) controls. ThsA N112A lacks SIR2-type NADase activity; ThsA R371A has an altered SLOG-motif, and ThsB E85Q lacks TIR-domain catalytic activity. All constructs expressed at OD 0.80. (*E*) Ion leakage assay in *Nb eds1*$^{-/-}$ leaves co-expressing different ThsA and ThsB combinations. Leaf discs were collected ~72 hpi, and ion measurements taken every 24 h for 3 days. High expression of AbTIR alone was shown to elicit NAD$^+$-depletion and cytotoxicity to a limited degree (see Fig. 1), therefore, the difference of samples co-expressed with inactive ThsA N112A controls was plotted to reveal ThsA-stimulation.

# SI 6

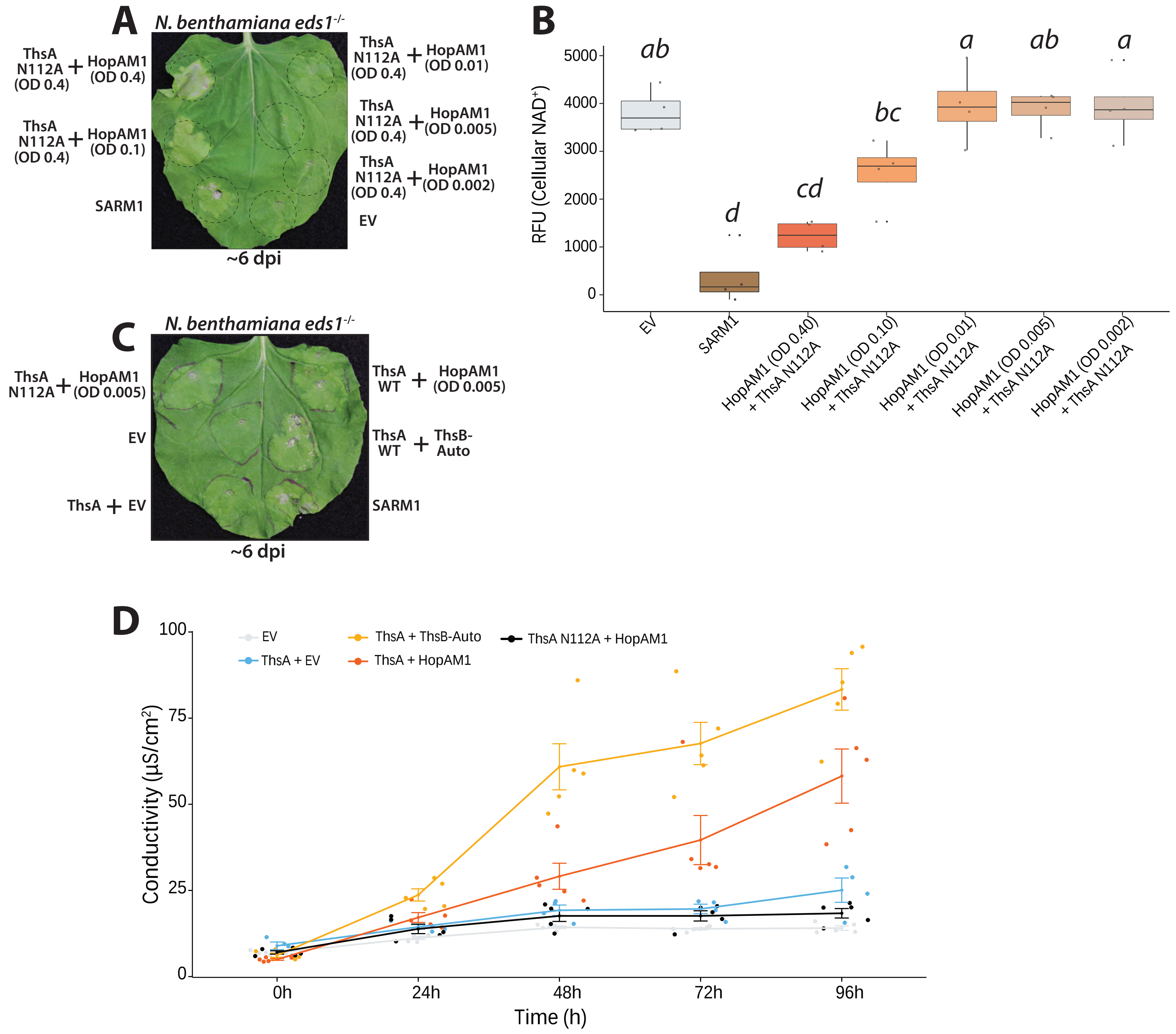


**SI 6**. HopAM1 causes dosage sensitive cell death independent of EDS1. (*A*) *Nb eds1*$^{-/-}$ leaves ~5 dpi with constructs expressing HopAM1 and ThsA N112A, or SARM1 and EV (35S:GFP) controls. ThsA N112A lacks SIR2-type NADase activity. The delivered dosage of HopAM1 was titrated as noted on the leaf. All other constructs were expressed at OD 0.80. (*B*) Fluorescent $NAD^+$-detection assay in *Nb eds1*$^{-/-}$ leaves from expressing different HopAM1 dosages along with ThsA N112A. Leaves harvested at 40 hpi. (*C*) *Nb eds1*$^{-/-}$ leaves co-expressing WT ThsA or N112A in combination with ThsB-Auto, HopAM1, or EV (35S:GFP, empty vector).

# SI7

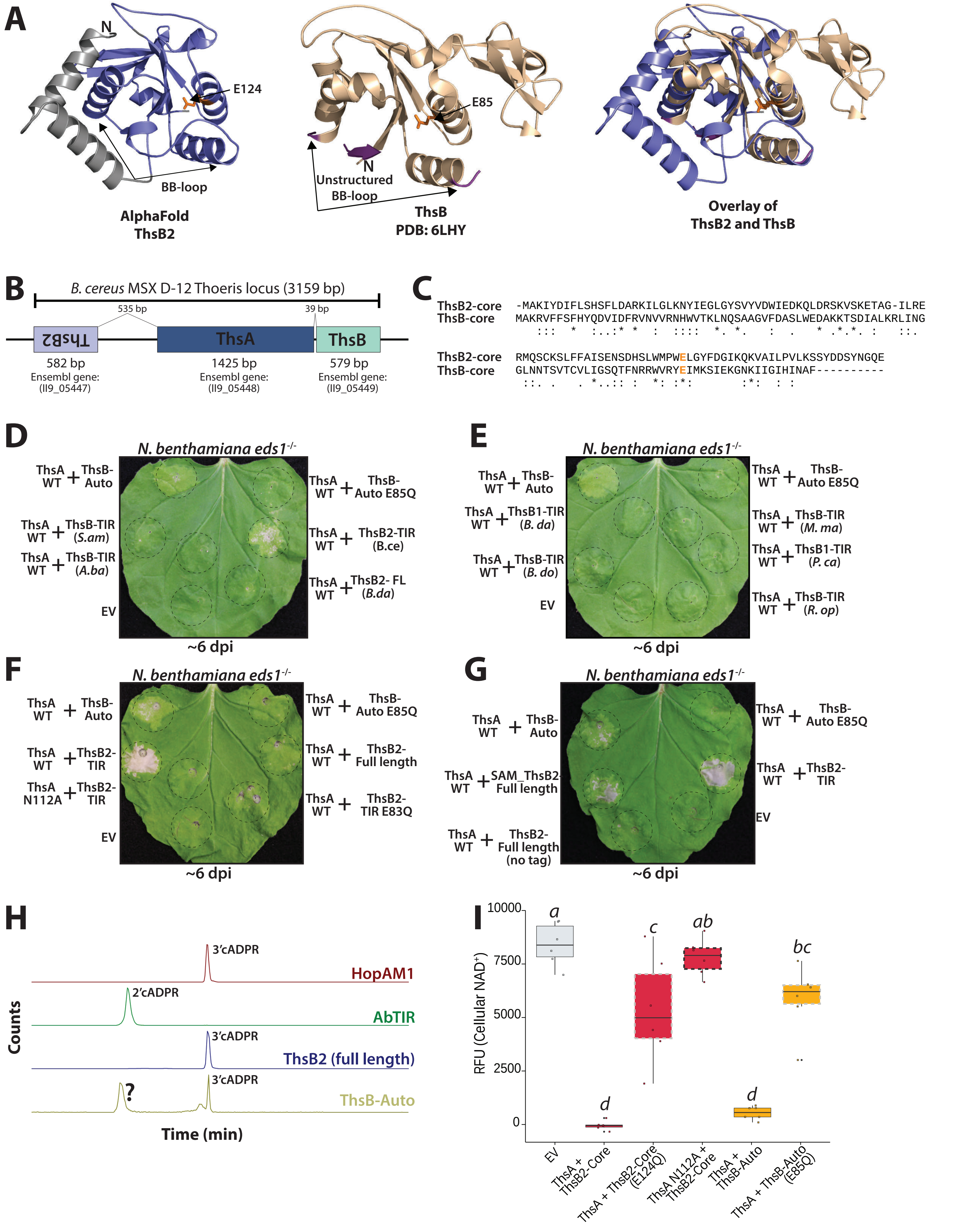
A
N
E124
BB-loop
AlphaFold ThsB2
E85
N
Unstructured BB-loop
ThsB PDB: 6LHY
Overlay of ThsB2 and ThsB
B
B. cereus MSX D-12 Thoeris locus (3159 bp)
535 bp
39 bp
ThsB2
ThsA
ThsB
582 bp
Ensembl gene: (II9_05447)
1425 bp
Ensembl gene: (II9_05448)
579 bp
Ensembl gene: (II9_05449)
C
ThsB2-core -MAKIYDIFLSHSFLDARKILGLKNYIEGLGYSVYVDWIEDKQLDRSKVSKETAG-ILRE
ThsB-core MAKRVFFSFHYQDVIDFRVNVVRNHWVTKLNQSAAGVFDASLWEDAKKTSDIALKRLING
ThsB2-core RMQSCKSLFFAISENSDHSLWMPWELGYFDGIKQKVAILPVLKSSYDDSYNGQE
ThsB-core GLNNTSVTCVLIGSQTFNRRWVRYEIMKSIEKGNKIIGIHINAF----------
D
N. benthamiana eds1-/-
ThsA WT + ThsB-Auto
ThsA WT + ThsB-TIR (S.am)
ThsA WT + ThsB-TIR (A.ba)
EV
ThsA WT + ThsB-Auto E85Q
ThsA WT + ThsB2-TIR (B.ce)
ThsA WT + ThsB2- FL (B.da)
~6 dpi
E
N. benthamiana eds1-/-
ThsA WT + ThsB-Auto
ThsA WT + ThsB1-TIR (B. da)
ThsA WT + ThsB-TIR (B. do)
EV
ThsA WT + ThsB-Auto E85Q
ThsA WT + ThsB-TIR (M. ma)
ThsA WT + ThsB1-TIR (P. ca)
ThsA WT + ThsB-TIR (R. op)
~6 dpi
F
N. benthamiana eds1-/-
ThsA WT + ThsB-Auto
ThsA WT + ThsB2-TIR
ThsA N112A + ThsB2-TIR
EV
ThsA WT + ThsB-Auto E85Q
ThsA WT + ThsB2-Full length
ThsA WT + ThsB2-TIR E83Q
~6 dpi
G
N. benthamiana eds1-/-
ThsA WT + ThsB-Auto
ThsA WT + SAM_ThsB2-Full length
ThsA WT + ThsB2-Full length (no tag)
ThsA WT + ThsB-Auto E85Q
ThsA WT + ThsB2-TIR
EV
~6 dpi
H
3'cADPR
HopAM1
2'cADPR
AbTIR
3'cADPR
ThsB2 (full length)
?
3'cADPR
ThsB-Auto
Counts
Time (min)
I
RFU (Cellular NAD+)
10000
7500
5000
2500
0
a
d
c
ab
d
bc
EV
ThsA + ThsB2-Core
ThsA + ThsB2-Core (E124Q)
ThsA N112A + ThsB2-Core
ThsA + ThsB-Auto
ThsA + ThsB-Auto (E85Q)

**SI 7**. The TIR-domain of a second ThsB from *B. cereus* MSX-D12 is auto-active and produces 3'-cADPR and stimulates ThsA; no other tested ThsB-allele stimulated ThsA. (*A*) AlphaFold structure prediction of the second ThsB (ThsB2) encoded by *B. cereus* MSX-D12 overlayed with the crystal structure of prototypical ThsB (PDB ID: 6LHY) of Ka *et al.* (*B*) Schematic of the *B. cereus* MSX-D12 Thoeris locus discovered by Doron *et al*. (*C*) ClustalW amino acid alignment of ThsB and ThsB2 reveals overall low residue identity. (*D-E*) *N. benthamiana eds1*$^{-/-}$ leaves expressing ThsA with ThsB-Auto or the ThsB-alleles screened in S2. (*F-G*) *N. benthamiana eds1*$^{-/-}$ leaves expressing ThsA with ThsB2-TIR (core TIR-domain) or full length ThsB2, as compared to ThsB-Auto. ThsB2-TIR alone does not cause cell death; catalytic E required to stimulate ThsA. (*H*) LC-MS analysis of *in vitro* NADase products from ThsB2, ThsB-Auto, and AbTIR or HopAM1. (*I*) Fluorescent $NAD^+$-detection assay in *Nb eds1*$^{-/-}$ leaves co-expressing ThsA WT (or ThsA N112A) with ThsB2-TIR or ThsB-Auto. E/Q substitutions in TIR-domain catalytic glutamate residue.

# SI8

| TIR | Organism | Known enzymatic product(s) | EDS1-HR? | EDS1-independent cytotoxicity? | ThsA-activation? |
|---|---|---|---|---|---|
| **HopAM1** | *Pseudomonas syringae (DC3000)* | 3'cADPR | X | ✓ | ✓ |
| **AbTIR** | *Acinetobacter baumanii* | 2'cADPR | X | ✓ | X |
| **TcpO-TIR** | *Methanobrevibacter olleyae* | v-cADPR (2'cADPR?) | X | ✓ | X |
| **ThsB-Auto** | *Bacillus cereus (MSX D-12)* | 3'cADPR, unknown isomer | X | X | ✓ |
| **ThsB2** | *Bacillus cereus (MSX D-12)* | 3'cADPR | X | X | ✓ |
| **BdTIR** | *Brachypodium distachyon* | 2'cADPR, 3'cADPR | ✓ | X* | ✓ |
| **RPP1** | *Arabidopsis thaliana* | 2'cADPR, 2',3'-cNMP, pRib-AMP, ADPr-ATP | ✓ | X | X |
| **SARM1** | *Homo sapiens* | cADPR, ADPR | X | ✓ | Not examined** (Ofir *et al*, 2021) |

**SI 8**. Summary of examined TIR-domains: reported enzymatic products and EDS1 / ThsA-signaling phenotypes. Orange background indicates TIR-proteins of prokaryotic origin; green indicates plant TIRs, and grey indicates human SARM1 TIR. *BdTIR can elicit *EDS1*-independent toxicity under specific contexts such as prolonged over-expression using Omega translational enhancers. **SARM1 not examined as cADPR was previously found to not stimulate ThsA (Ofir *et al*., 2021).
